# YAP and TAZ are transcriptional co-activators of AP-1 proteins and STAT3 during breast cellular transformation

**DOI:** 10.1101/2021.02.18.431832

**Authors:** Lizhi He, Henry Pratt, Fengxiang Wei, Mingshi Gao, Zhiping Weng, Kevin Struhl

**Author notes:** To whom correspondence should be addressed at and. These authors contributed equally (Lizhi He, Henry Pratt, Fengxiang Wei).

## Abstract

The YAP and TAZ paralogues are transcriptional co-activators recruited to target sites, primarily by TEAD proteins. Here, we show that YAP and TAZ are also recruited by JUNB and STAT3, key factors that mediate an epigenetic switch linking inflammation to cellular transformation. YAP and TAZ directly interact with JUNB and STAT3 via a WW domain important for transformation, co-occupy many target sites *in vivo* via AP-1 and (to a lesser extent) STAT3 sequence motifs, and stimulate transcriptional activation by AP-1 proteins. A few target sites are YAP- or TAZ-specific, and they are associated with different sequence motifs and gene classes. YAP/TAZ, JUNB, and STAT3 directly regulate a common set of target genes that overlap, but are distinct from, those regulated by YAP/TAZ and TEADs. The set of genes regulated by YAP/TAZ, STAT3, and JUNB is associated with poor survival in breast cancer patients with the triple-negative form of the disease.

## INTRODUCTION

The Hippo signal transduction pathway plays critical roles in development, homeostasis and tumor progression (Piccolo et al., 2014; Varelas, 2014; Yu et al., 2015; Totaro et al., 2018; Moya and Halder, 2019). Internal and external signals relayed through the Hippo pathway converge on YAP and TAZ, paralogous proteins with 63% sequence similarity that are the major effectors of this pathway (Piccolo et al., 2014; Totaro et al., 2018; Ma et al., 2019). Once activated, YAP/TAZ translocate into nucleus and act as transcriptional co-activators, most notably by interacting with the TEAD family of DNA-binding transcription factors (Piccolo et al., 2014; Totaro et al., 2018; Ma et al., 2019; Moya and Halder, 2019). Deregulation of YAP/TAZ activity is frequently observed in various human cancers, contributing to cancer initiation, progression, and metastasis (Johnson and Halder, 2014; Yu et al., 2015; Zanconato et al., 2016). YAP/TAZ activation is also linked to chemo-resistance in cancer therapy (Johnson and Halder, 2014; Yu et al., 2015; Zanconato et al., 2016). However, the transcriptional mechanisms and regulatory circuits mediated by YAP/TAZ in cancers, especially those beyond their association with TEAD1-4, are not well understood.

In addition to directly interacting with the TEAD family of DNA-binding proteins, YAP/TAZ can function through other mechanisms. Via its interaction with TEAD proteins, YAP/TAZ can synergize with JUN/FOS at composite regulatory elements containing both TEAD and AP-1 motifs (Zanconato et al., 2015). YAP can form a functional complex with β-catenin and TBX5 in cancer cells (Rosenbluh et al., 2012), YAP/TAZ can integrate into a SMAD-OCT4 complex in embryonic stem cells (Beyer et al., 2013), and a YAP-p73 complex plays a role in the DNA damage response (Beyer et al., 2013). In addition, YAP/TAZ directly interacts with the general co-activator BRD4 to increase RNA polymerase II recruitment and transcription of target genes (Zanconato et al., 2018), and YAP/TAZ activity is inhibited by a direct interaction with the SWI/SNF complex via the ARID1A subunit (Chang et al., 2018).

In previous work, we described an epigenetic switch that transforms breast cells via the inflammatory regulatory network controlled by the joint action of NF-κB, STAT3, and AP-1 transcription factors (Iliopoulos et al., 2009; Iliopoulos et al., 2010; Ji et al., 2018; Ji et al., 2019). Here, we show that YAP and TAZ are important for cellular transformation in this breast cell transformation model. We demonstrate that YAP and TAZ directly interact with STAT3 and JUNB and act as transcriptional co-activators that stimulate expression of target genes. These target genes overlap with, but are distinct from, the target genes in which YAP/TAZ is recruited by TEAD proteins. The YAP/TAZ target genes mediated by STAT3/AP-1, but not TEAD proteins, are associated with poor survival prognosis in breast cancer patients with the triple-negative form of the disease.

## RESULTS

### YAP and TAZ are important for STAT3 and AP-1 activity during breast epithelial cell transformation

We examined the role of YAP and TAZ in breast cellular transformation by using CRISPR to knock out these genes in the context of our Src-inducible model. Knockout of YAP or TAZ or both (Figure 1A) reduces growth under conditions of low attachment (Figure 1B) and inhibits colony formation in soft agar (Figure 1C). In contrast, there is no effect on cell proliferation under standard conditions of high attachment (Figure 1D). Similar results are observed when YAP or TAZ expression is knocked down by siRNA (Figure 1E-G). Thus, YAP and TAZ are important for cellular transformation in this model.

**Figure 1.**
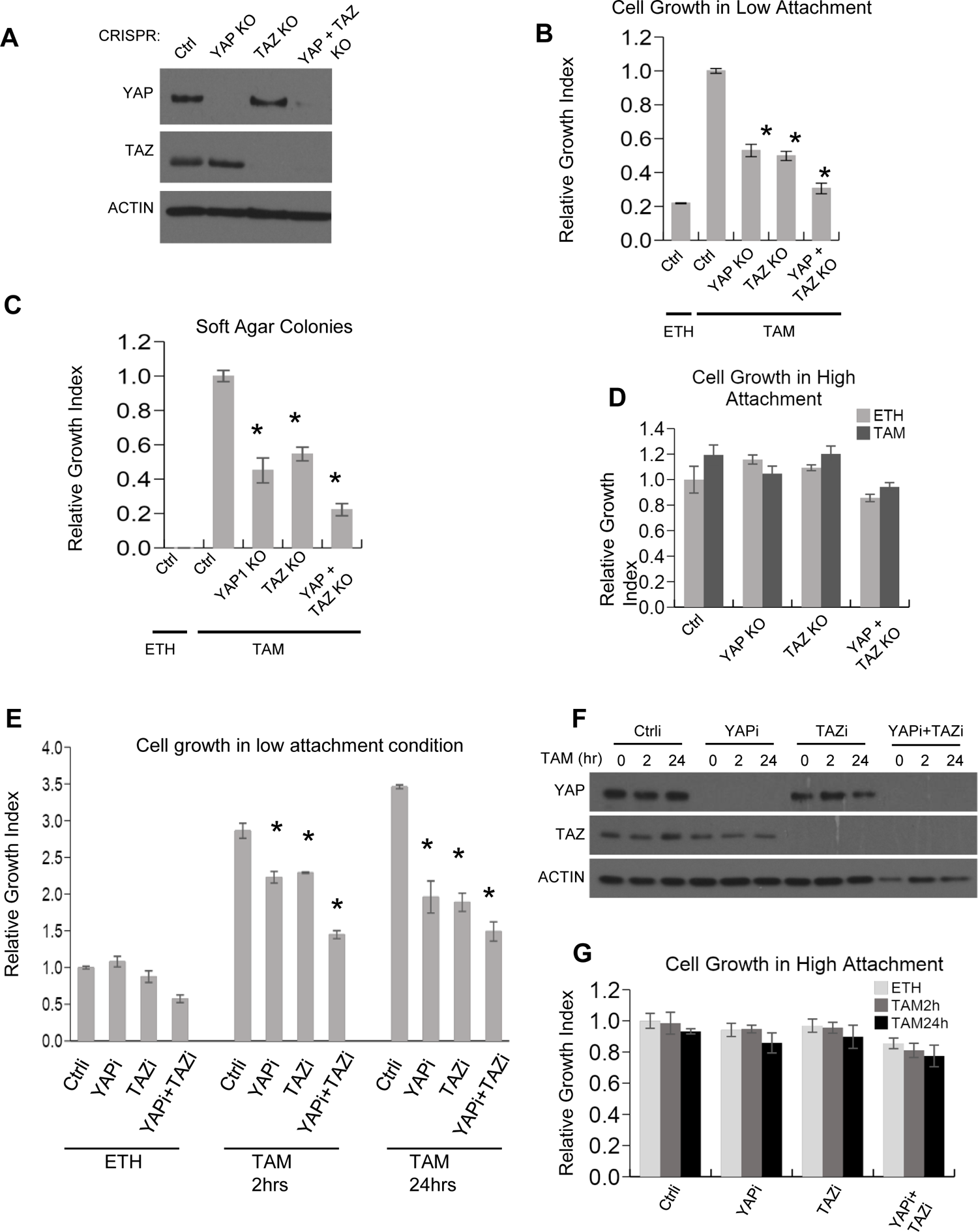
YAP and TAZ facilitate transformation. (A) Western blot for YAP, TAZ, and Actin levels in the indicated CRISPR-mediated knockout (KO) strains and the parental cell line (Ctrl). (B) Relative growth in low attachment conditions of the indicated cell lines in non-transformed (ETH) and transformed conditions (TAM). Measurements are relative to transformed cells with the CRISPR control, which is defined as 1.0. (C) Number of colonies growing in soft agar in the indicated cell lines. (D) Relative growth of the indicated cell lines in standard high attachment conditions. (E) Relative growth in low attachment conditions, (F) YAP and TAZ protein levels, and (G) relative growth in high attachment conditions of cells subjected to siRNA-mediated knockdown of YAP and/or TAZ (YAPi and/or TAZi) of cells induced by TAM addition for the indicated times. Error bars indicate ± SD of 3 replicates and * indicates values significantly different (*p* < 0.05) from the control.

An inflammatory regulatory network mediated by the joint action of NF-κB, STAT3, and AP-1 factors at common target sites is critical for transformation in our inducible model, and the network is involved in many human cancers (Ji et al., 2019). Using siRNA-mediated knockdowns, we examined whether YAP and TAZ are important for the increased activity of these transcription factors during transformation. Transient knockdown of YAP and/or TAZ, but not JUNB, strongly decreases STAT3 phosphorylation at Tyr705 (STAT3-p, a marker of STAT3 activation) during transformation (Figure 2A), but it does not affect overall levels of STAT3 protein (Figure 2A) or mRNA (Figure 2-figure supplement 1). Consistent with this observation, YAP and TAZ are important for induction of IL6 transcription and secretion during transformation (Figure 2B). In addition, depletion of YAP or TAZ (and also STAT3 or JUNB) inhibits transcriptional activation mediated by AP-1 factors (Figure 2C). In contrast, YAP or TAZ depletion causes a slight increase in NF-κB activation (Figure 2D), perhaps reflecting competition between the Hippo and NF-κB signaling pathways. The effects of YAP and TAZ on AP-1 and STAT3 activity during transformation are not due to changes in YAP or TAZ protein levels in the nucleus (Figure 3A).

**Figure 2.**
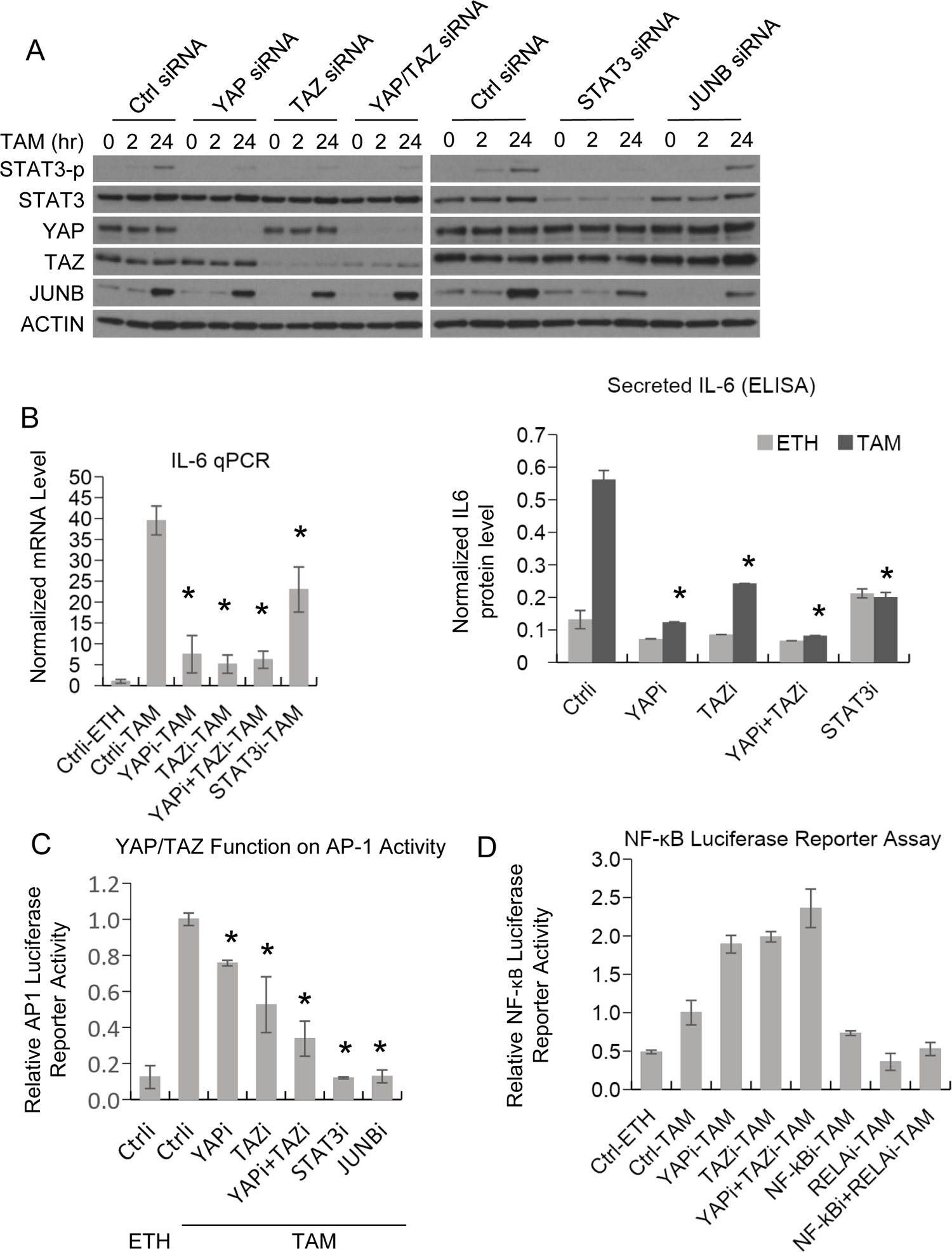
YAP and TAZ regulate STAT3 and JUNB activities during transformation. (A) Western blot for the indicated proteins (STAT3-p, is the form phosphorylated at Tyr705) in cells treated with the indicated siRNAs and with TAM for the indicated times. (B) Normalized IL-6 mRNA (left) and secreted IL-6 (right) levels in the indicated cells and conditions. (C) Relative AP-1-dependent transcriptional activity in the indicated cells and conditions. (D) Relative NF-κB-dependent transcriptional activity. Error bars indicate ± SD of 3 replicates and * indicates values significantly different (*p* < 0.05) from the control.

**Figure 3.**
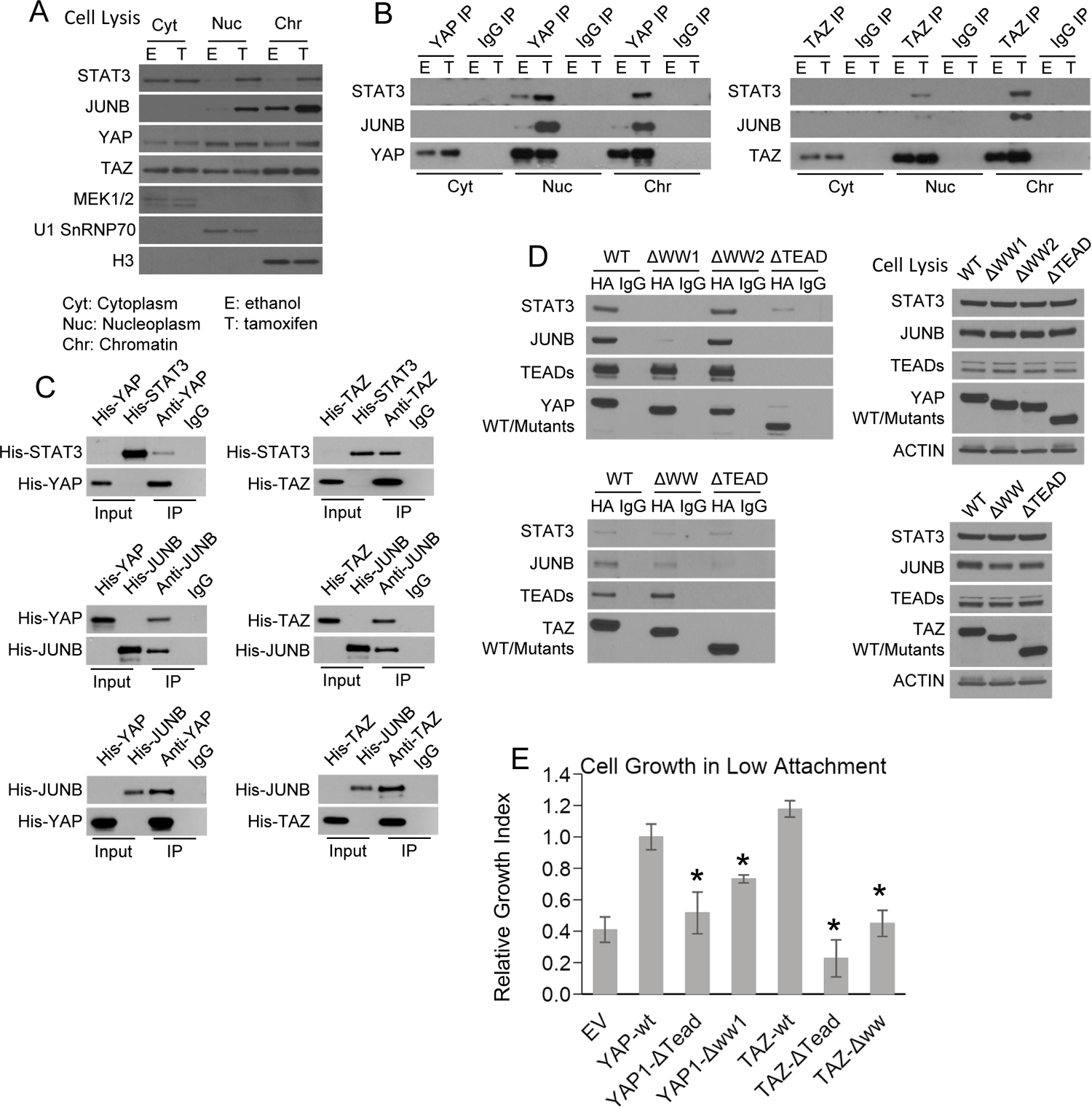
YAP and TAZ, via WW domains, directly interact with STAT3 and JUNB. (A) Levels of YAP, TAZ, STAT3 and JUNB in the indicated fractions in non-transformed (E) and transformed (T) conditions: MEK1/2, cytoplasm marker; U1 SnRNAP70, nucleoplasm marker; H3, chromatin marker. (B) Co-immunoprecipitation in cellular fractions. Western blot of the indicated proteins upon immunoprecipitation (IP) with antibodies against the indicated proteins or the IgG control. (C) Co-immunoprecipitation with *E. coli*-generated His-tagged recombinant proteins. Western blot of the input and immunoprecipitated (IP) proteins with antibodies against the indicated proteins. The input sample contained 10% of the amount used for recombinant proteins used for the co-immunoprecipitation. (D) Left panels are Western blots of the indicated proteins upon immunoprecipitation with the indicated HA-tagged YAP (top) or TAZ (bottom) derivatives or IgG control. Right panels are Western blot of cell extracts prior to immunoprecipitation. (E) Relative growth in conditions of low attachment in cells overexpressing the indicated proteins or empty vector (EV) control in parental MCF-10A cells (lacking ER-Src). Error bars indicate ± SD of 3 replicates and * indicates values significantly different (*p* < 0.05) from the control.

### YAP and TAZ directly interact with JUNB and STAT3 during transformation

The effects of YAP and TAZ knockout on STAT3 and AP-1 activity could be direct, or they could be an indirect consequence of inhibiting transformation. Among the many members of the AP-1 transcription factor family, expression of JUNB is most strongly induced during transformation (Figure 3-figure supplement 1A), leading us to study JUNB in more detail. We addressed the possibility of a direct effect by co-immunoprecipitation (co-IP) experiments in cytoplasmic, nucleoplasm, and chromatin fractions from non-transformed and transformed cells. YAP and TAZ proteins are observed in all three cellular fractions, and their levels are unchanged upon tamoxifen treatment (Figure 3A and Figure 3-figure supplement 1B). This indicates that the Hippo pathway is active in this cell line prior to transformation and is not altered during the transformation process. In the nucleoplasm and chromatin (but not cytoplasm) fractions, YAP and TAZ each co-immunoprecipitate with STAT3 and JUNB (Figure 3B). In general, co-IP is more efficient in transformed cells. We confirmed direct pairwise YAP-STAT3, TAZ-STAT3, YAP-JUNB, and TAZ-JUNB interactions by performing co-IP experiments using histidine-tagged recombinant proteins generated in *E. coli* (Figure 3C).

### WW domains of YAP and TAZ are important for interacting with STAT3 and JUNB and for transformation

YAP and TAZ have a TEAD domain that interacts with TEAD1-4, and they contain WW domains (two for YAP and one for TAZ) that mediate interactions with a variety of other proteins. To map the regions of YAP and TAZ required for interacting with STAT3 and JUNB, we performed co-IP experiments in cells co-expressing YAP or TAZ derivatives lacking one or both of these WW domains along with FLAG-tagged STAT3 or JUNB (Figure 3D). The WW1 domain of YAP and the WW domain of TAZ are critical for the interaction with STAT3 and JUNB, whereas removal of the YAP WW2 domain has no effect on these interactions. Removal of any WW domain has no effect on the interaction with the TEAD proteins, whereas removal of the TEAD domain abolishes the interaction with the TEAD proteins. In addition, the TEAD domains of both YAP and TAZ are critical for the interaction with JUNB, but they contribute only partially to the interaction with STAT3 (Figure 3D). When overexpressed in parental MCF-10A cells (i.e., lacking the ER-Src protein), wild-type YAP or TAZ increase the level of transformation, whereas derivatives lacking YAP-WW1 or TAZ-WW do not (Figure 3E). Like the wild-type proteins, none of these derivatives have a significant effect on cell proliferation under conditions of high attachment (Figure 3-figure supplement 1C). Thus, the WW1 domain in YAP and the WW domain in TAZ, which are critical for interaction with JUNB and STAT3 but not TEAD proteins, are important for transformation. These observations indicate that YAP and TAZ interactions with TEAD proteins are not sufficient for transformation, and they suggest that the YAP and TAZ interactions with JUNB and/or STAT3 are important for transformation.

### YAP and TAZ have highly similar, but not identical, genomic binding profiles

Using protein-specific antibodies, we performed ChIP-seq to identify the genomic target sites of YAP, TAZ, STAT3, JUNB, and TEAD in both non-transformed and transformed cells. In general, there is excellent agreement in genome-wide signal profiles between pairs of biological replicates for each factor (Table S1A), and the binding regions of the three sequence-specific transcription factors—STAT3, JUNB, and TEAD—are strongly enriched in their respective motifs (Figure 4-figure supplement 1). YAP, TAZ, JUNB, and TEAD each have similar numbers of peaks in the non-transformed and transformed conditions, while STAT3 binding sites increase five-fold upon transformation (Table S1B). As expected from their being paralogues, YAP and TAZ have very similar binding profiles in both the transformed and non-transformed states (Figure 4A; overall Pearson correlation coefficient ∼0.7). For most subsequent analyses, we will consider these shared YAP/TAZ sites together.

**Figure 4.**
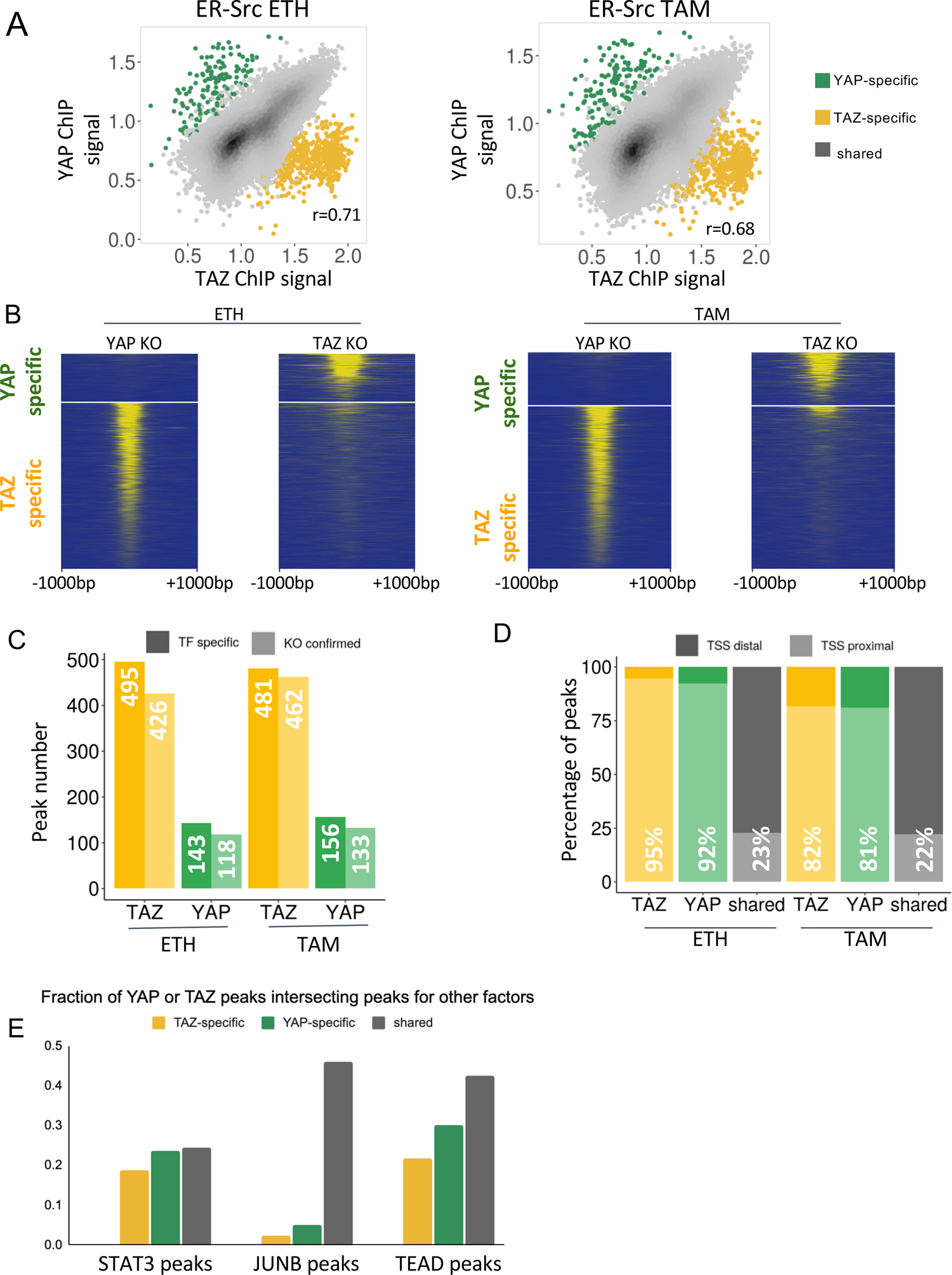
YAP and TAZ have highly similar binding profiles, but a subset of binding sites are unique to each factor. (A) Correlation (r ∼ 0.7) of YAP and TAZ binding signals in non-transformed (ETH) and transformed (TAM) cells. Putative YAP- and TAZ-specific sites are indicated, respectively, as green and yellow. (B) YAP and TAZ binding signals in YAP- or TAZ-knockout cell lines using an antibody that recognizes both proteins. (C) Number of putative YAP- and TAZ-specific sites (dark colors) confirmed in cells deleted for the indicated factor (lighter colors). (D) Percentage of YAP-specific, TAZ-specific, and shared YAP/TAZ sites that are located proximal (light colors) or distal (darker colors) to the transcription start site (TSS). (E) Percentages of TAZ-specific (yellow), YAP-specific (green), and YAP/TAZ shared (gray) sites intersecting with STAT3, JUNB, and TEAD sites in transformed cells. TAZ- and YAP-specific sites are significantly less likely than shared sites to intersect JUNB and TEAD sites (Chi-square *p*-values YAP/JUNB = 3.2×10^-24^, TAZ/JUNB = 2.1×10^-81^, YAP/TEAD = 0.002, TAZ/TEAD = 9.7×10^-20^).

### YAP- and TAZ-specific sites differ in their associated motifs and biological functions from share YAP/TAZ sites

Unexpectedly, ∼575 target sites (∼2% of YAP/TAZ sites) appear specific for either YAP or TAZ (Figure 4A). To test whether these apparent YAP- or TAZ-specific sites are false positives or false negatives, we knocked out YAP and TAZ separately in ER-Src cells and then performed ChIP-seq experiments using an antibody that recognizes both proteins (Figure 4B). When compared to the parental cell line, a YAP-specific site should show reduced binding only in the YAP-deletion line, and a TAZ-specific site should show reduced binding only in the TAZ-deletion line. In this manner, we confirmed ∼450 TAZ-specific sites and ∼125 YAP-specific sites (Figure 4C).

Several characteristics of these YAP- and TAZ-specific sites suggest they serve biologically distinct functions from shared sites. First, > 80% of YAP- and TAZ-specific sites are located within 150 base pairs upstream of a GENCODE-annotated transcription start site (TSS), but < 25% of shared YAP/TAZ sites are so localized (Figure 4D). Second, YAP- and TAZ-specific sites are much less likely to be located within JUNB target regions (Chi-square *p*-value=3.2×10^-24^ for YAP, 2.1×10^-81^ for TAZ) and TEAD (Chi-square *p*-value=0.002 for YAP, 9.7×10^-20^ for TAZ) than shared YAP/TAZ sites (Figure 4E). Third, GO analysis of the genes whose promoters are near YAP- and TAZ-specific sites identify translation, rRNA processing, and mitochondrial gene expression as overrepresented gene categories for TAZ-specific sites, and transcriptional regulation and chromatin remodeling as categories overrepresented for YAP-specific sites (Table S2). Fourth, the TAZ-specific peaks are enriched in unique motifs (Figure 4-figure supplement 1 and see below).

### YAP and TAZ co-occupy genomic target sites with STAT3, JUNB, and TEAD proteins

We compared the YAP/TAZ binding data with that of STAT3, JUNB, and TEAD proteins (Figure 5A, Figure 5-figure supplement 1A). As expected, about 90% of the TEAD target sites are also bound by YAP or TAZ. Meanwhile, 55% of STAT3 and 42% of JUNB target sites are bound by YAP/TAZ, and reciprocally, 55% of YAP/TAZ peaks are bound by STAT3 or JUNB (Figure 5A). These co-occurrences are highly significant because a control set of comparable DNase hypersensitivity sites (DHSs) that are not bound YAP/TAZ show far lower co-occurrence of these transcription factors (Figure 5B). TEAD, STAT3, or JUNB are present, respectively in 3%, 9%, and 19% of control DHSs compared with 40%, 23%, and 42% of YAP/TAZ sites (Chi-square *p*-value <10^-100^ for all comparisons; Figure 5B). Together with the physical interactions among these proteins (Figure 3B, C), these observations strongly suggest that YAP/TAZ co-occupy genomic sites with STAT3 and JUNB.

**Figure 5.**
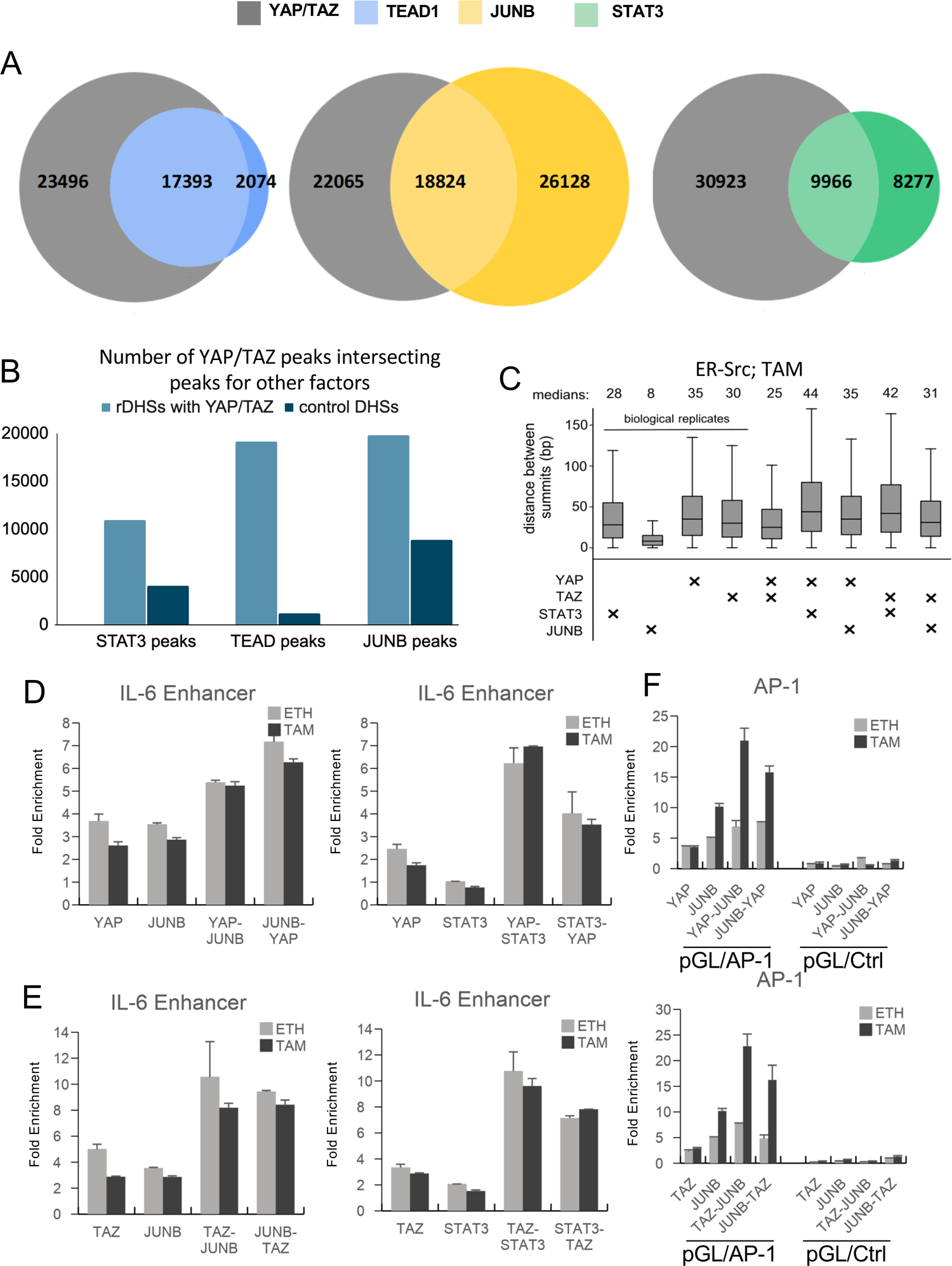
YAP and TAZ co-occupy sites with JUNB and STAT3. (A) Venn Diagrams for the intersection of the indicated factor binding sites in transformed cells. (B) Number of DNase hypersensitivity sites containing a YAP/TAZ peak (light blue) and matched control DHSs (dark blue) intersecting STAT3, JUNB, and TEAD sites; YAP/TAZ-associated DHSs are significantly more likely to intersect peaks for the three factors (Chi-square *p*-values <10^-100^ for all comparisons). (C) Distance between peak summits for biological replicates or combinations of the indicated factors in transformed cells. (D) Fold-enrichments of individual and sequential ChIP at the IL-6 enhancer in untransformed (ETH) and transformed (TAM) cells with YAP and either JUNB or STAT3 performed in the indicated order. (E) Fold-enrichments of individual and sequential ChIP at the IL-6 enhancer with TAZ and either JUNB or STAT3 performed in the indicated order. (F) Fold-enrichments of individual and sequential ChIP of the indicated proteins on a plasmid containing either 6 AP-1 motifs or a control lacking these motifs (Ctrl). Error bars indicate ± SD of 3 replicates and * indicates values significantly different (*p* < 0.05) from the control.

We further investigated protein co-occupancy by performing pairwise analysis of ChIP-seq peak summits. For biological replicates of a given protein in transformed cells, the median distances between corresponding peak summits are between 8-35 bp, and this serves as a control. YAP and TAZ peak summits are separated by 25 bp, indicating that these two proteins occupy the same sequences, although not necessarily at the same time (Figure 5C). Interestingly, the median distances between pairwise combinations of YAP or TAZ with STAT3 or JUNB range between 31-44 bps (Figure 5C), indicating that YAP and TAZ binding occurs very close to the location of STAT3 and JUNB. Similar results are observed in non-transformed cells (Figure 5-figure supplement 1B).

We demonstrated co-occupancy of YAP and TAZ with STAT3 and JUNB by sequential ChIP experiments at selected loci. At the IL-6 enhancer, sequential ChIP in either order yields increased fold-enrichments of YAP (Figure 5D) or TAZ (Figure 5E) with JUNB or STAT3 in non-transformed and transformed cells; similar results are observed at the SNX24 locus (Figure 5-figure supplement 1C). At the MYC locus, increased fold-enrichment is generally seen only in one direction (JUNB or STAT3 first, with the exception of TAZ and STAT3; Figure 5-figure supplement 1C-D), suggesting that most of the YAP and TAZ binding is not mediated by JUNB or STAT3, although they can co-occupy the locus. In addition, sequential ChIP (in both orders) of JUNB and either YAP or TAZ on a transfected plasmid-borne locus containing six AP-1 motif sites shows increased fold-enrichment in transformed cells, while this does not occur on a control plasmid locus lacking the AP-1 sites (Figure 5F).

### YAP and TAZ are recruited via AP-1 and, to a lesser extent, STAT3 motifs

Transcriptional co-activators such as YAP and TAZ are ultimately recruited to genomic sites by sequence-specific DNA-binding proteins. Thus, motifs enriched in YAP and TAZ binding sites suggest which proteins mediate the recruitment. In ER-Src cells, JUNB binds primarily to AP-1 sequence motifs, indicative of a direct protein-DNA interaction, whereas STAT3 binding sites contain either STAT3 or AP-1 motifs, indicating that STAT3 can bind directly to DNA or indirectly via interactions with AP-1 proteins (Ji et al., 2019). To determine which motifs are enriched at YAP and TAZ sites, we compared the frequency of sequence motifs from the HOCOMOCO and JASPAR catalogs within YAP/TAZ target sites against comparable DNase hypersensitive sites that are not bound by YAP/TAZ.

As expected from YAP/TAZ being a co-activator of TEAD proteins, ∼30% of YAP/TAZ binding loci contain TEAD motifs (Figure 6A), and the TEAD motif is the most significantly enriched motif within YAP/TAZ sites (Figure 6B). Interestingly, and in accord with the direct interaction of JUNB and YAP/TAZ, a comparable number of YAP/TAZ target sites contain AP-1 motifs (Figure 6A). However, the fold-enrichment of AP-1 motifs at YAP/TAZ sites vs. control loci is only modest (Figure 6B), presumably due to AP-1 factors being involved in a wide range of pathways. YAP and TAZ target sites also contain STAT3 motifs, albeit at much lower frequency (13%) and minimal enrichment over control loci. Importantly, all the TEAD, AP-1, and STAT3 motifs are centered around YAP/TAZ peak summits (Figure 6C), as expected for direct recruitment of YAP/TAZ by these DNA-binding transcription factors. In accord with previous observations (Zanconato et al., 2015), ∼6% of YAP/TAZ target sites contain both AP-1 and TEAD motifs (Figure 6D), a higher frequency than expected by chance (Figure 6E). In contrast, NF-κB motifs are not enriched or centered at YAP/TAZ target sites (Figure 6A-C). Taken together with the direct protein-protein interactions, these results strong suggest that, in addition to TEADs, AP-1 proteins and (to a lesser extent) STAT3 can directly recruit YAP/TAZ to genomic sites.

**Figure 6.**
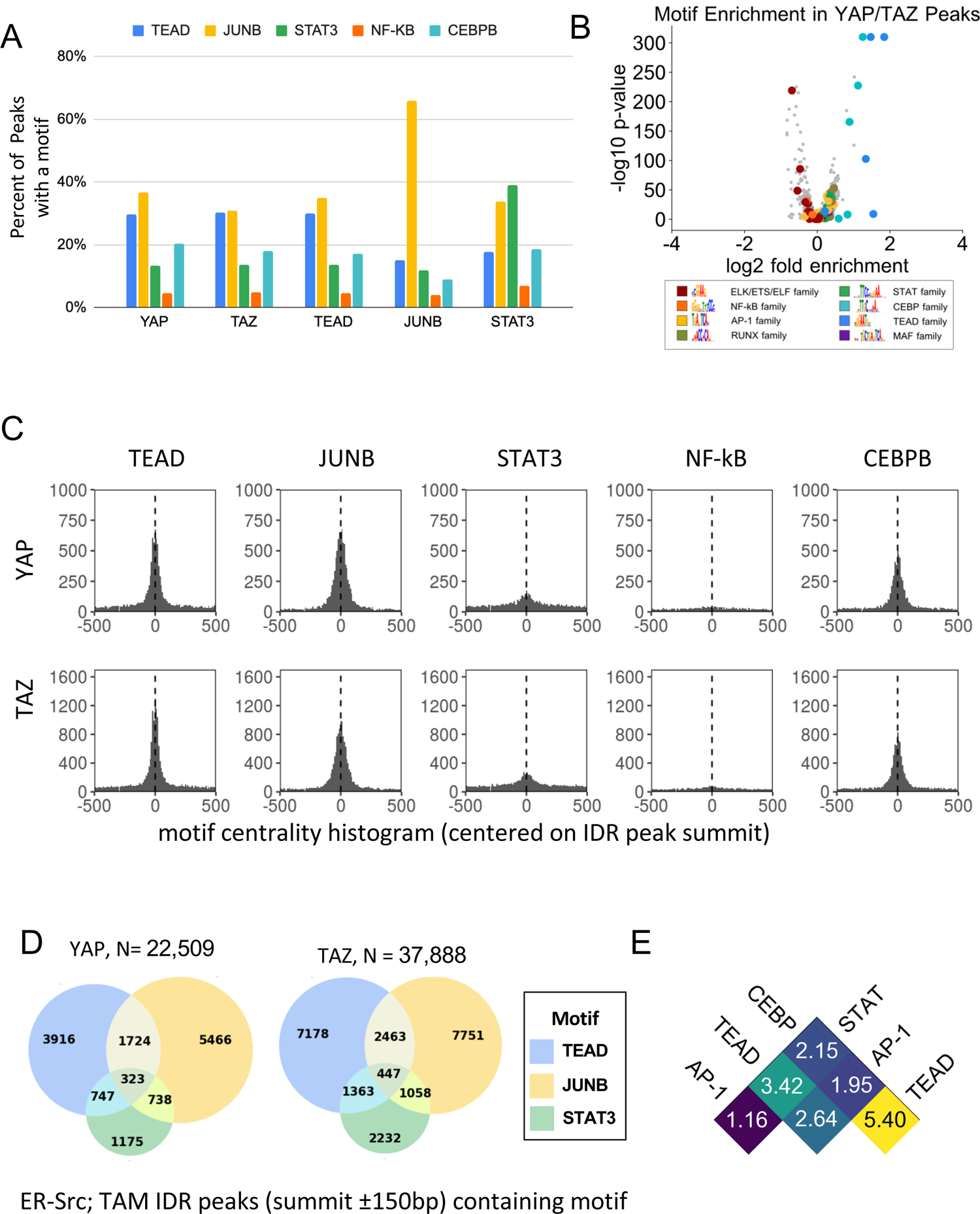
YAP/TAZ are recruited by TEAD, AP-1, CEBP proteins, and (to a lesser extent) STAT3. (A) Percent of binding regions for the indicated proteins that contain a given DNA sequence motif (different colors). (B) Enrichment and *p-*values for various motifs within YAP/TAZ target sites in transformed cells as compared with control sites having similar DNase-seq signal profiles. (C) Histogram of all motif locations for TEAD, JUNB, STAT3, NF-κB, and CEBPB occurring within 500 bp of a YAP (top) or TAZ (bottom) peak summit (defined as position 0). (D) Overlap between YAP and TAZ peaks containing the indicated motifs. (E) Fold-enrichment of the indicated pairwise combinations of motif in YAP/TAZ target sites relative to control sites.

### CEBP motifs are enriched at YAP/TAZ target sites, and TAZ-specific sites are enriched for ETS motifs and depleted for AP-1 motifs

Unexpectedly, the motif recognized by the CEBP family of transcription factors occurs in ∼25% of YAP/TAZ sites with an enrichment over control sites that is second-most behind that of TEAD motifs (Figure 6B). CEBP motifs also occur very close to YAP/TAZ peak summits (Figure 6C), suggesting that one or more CEBP proteins can recruit YAP/TAZ to target sites. In addition, CEBP motifs are strongly enriched within TEAD binding regions (Figure 4-figure supplement 1), and CEBP and TEAD motifs co-occur at YAP/TAZ sites at more than five times the rate that they co-occur in control regions (Figure 6E). Although CEBP proteins have not been studied in the ER-Src model, it is noteworthy that CEBP_β_ ranks fourth among transcription factors predicted to be important for transformation (just behind JUNB, STAT3, and FOSL1), and siRNA-mediated depletion of CEBP_β_ reduces the level of transformation (Ji et al., 2018). Lastly, a minority of YAP/TAZ binding sites lack AP-1, STAT3, TEAD, or CEBP motifs, suggesting that other DNA-binding proteins can also recruit these co-activators.

Analysis of ∼450 TAZ-specific sites (Figure 4) indicates that the aforementioned motifs are not significantly enriched (Figure 4-figure supplement 1, lower-right panel; there are insufficient YAP-specific sites to perform this analysis). Instead, motifs for ETS family proteins are strongly enriched and the AP-1 motif is significantly depleted at TAZ-specific sites compared to control sites. This observation strongly supports the idea that TAZ-specific sites represent a biologically distinct subset from YAP/TAZ shared sites that differ with respect to the mechanism of co-activator recruitment and the genes that are affected.

### YAP and TAZ co-occupy sites with JUNB and STAT3 in a triple-negative breast cancer cell line

To provide independent support of our results, we performed ChIP-seq in a triple-negative breast cancer cell line (MDA-MB-231). Binding sites for YAP, TAZ, STAT3, JUNB, and TEAD (Table S1B) show considerable overlap with binding sites in ER-Src cells, although many sites appear cell-line specific (Figure 6-figure supplement 1). In MDA-MB-231 cells, YAP and TAZ have similar binding sites (Figure 6-figure supplement 2A) that frequently coincide with TEAD, AP-1, and STAT3 binding sites (Figure 6-figure supplement 2B, C). The percentage of YAP/TAZ binding sites associated with AP-1, TEAD, and STAT3 motifs are comparable in both cell lines (compare Figure 6A with Figure 6-figure supplement 3A, B), and roughly half of the YAP/TAZ target sites lack these motifs. These motifs are strongly enriched very near YAP/TAZ peak summits (Figure 6-figure supplement 3C), and similar ratios of motif co-occurrences within YAP/TAZ sites are observed in both cell lines (Figure 6-figure supplement 3D). Lastly, the 267 TAZ-specific sites in MDA-MB-231 cells are enriched for ELF/ETS-family motifs (Figure 6-figure supplement 3E). Thus, the binding profiles of these factors, individually and in combination, are very similar in both cell lines.

Interestingly, there are differences between the two cell lines with respect to how YAP/TAZ is recruited to target loci. At YAP/TAZ target sites, the enrichment of CEBP motifs in ER-Src cells is not observed in MDA-MB-231 cells (Figure 6-figure supplement 3A, B). Conversely, motifs recognized by the RUNX tumor suppressor are significantly enriched MDA-MB-231 cells (Figure 6-figure supplement 3B), but not in ER-Src cells. Thus, the putative recruitment of YAP/TAZ by CEBP proteins and RUNX appears to be cell-type-dependent. This specificity could be due to differences in levels or post-translational modifications of the recruiting proteins, and it is likely to cause cell-type-specific differences in gene expression.

### Different classes of YAP/TAZ sites are associated with different categories of genes

We asked whether subsets of YAP/TAZ binding sites in different cell lines or recruited by different transcription factors regulate biologically distinct subsets of genes. Of the 14,851 genes having a YAP/TAZ binding site within 2 kb of a TSS, 42% have a YAP/TAZ site in both cell lines, while 40% and 18% are specific to MDA-MB-231 and ER-Src, respectively (Figure 6-figure supplement 4A). Of 9,690 target genes with a proximal YAP/TAZ site in ER-Src cells, a core set of 1,562 were associated with binding sites for all four of YAP, TAZ, JUNB, and STAT3 (Figure 6-figure supplement 4B), and 6,806 (70%) were associated with a YAP/TAZ site with a candidate recruiting motif (Figure 6-figure supplement 4C). GO analysis for YAP/TAZ target genes specific for ER-Src vs. MDA-MB-231-yielded no significant terms at a false discovery rate (FDR) < 5%. However, genes targeted by YAP/TAZ sites do differ in their GO enrichment terms based on the presumed recruiting motif. Sites with an AP-1, CEBP, or TEAD motif are enriched for ontology terms related, respectively, to development, inflammation, and actin cytoskeletal organization at FDR < 5% (Figure 6-figure supplement 4D, Table S3), Terms related to cell adhesion and signal transduction are shared between these three classes, albeit at mild enrichment. No significantly enriched GO terms are observed for YAP/TAZ sites with STAT3 motifs (Table S3).

### A core set of genes is regulated by YAP, TAZ, STAT3, and JUNB

To identify genes regulated by YAP, TAZ, JUNB, STAT3, or TEAD proteins, we individually depleted these factors by siRNA-mediated knockdown in tamoxifen-treated ER-Src cells. RNA-seq analysis identifies between 1,000 and 4,000 differentially expressed genes for each condition as compared with a control siRNA (FDR < 5%; Figure 7A-B). Roughly equal number of genes show increased or decreased expression with the directionality of differential expression nearly always preserved among factors (Figure 7A-C). In accord with previous results (Ji et al., 2018), there is significant overlap between the differentially expressed genes identified for each factor, with a majority of YAP- or TAZ-affected genes being affected by at least one other factor (Figure 7A-C). We identified a core set of genes that are affected by all of YAP, TAZ, JUNB, and STAT3 (Figure 7A). Another core set of 292 genes, of which 149 decrease in expression and 143 increase, are affected by YAP, TAZ, and TEADs (Figure 7B). Notably, binding of these transcription factors is significantly more frequent at promoters of these core genes than at randomly generated control sets of protein-coding genes (Figure 7D; *p*-value < 0.01 for all factors).

**Figure 7.**
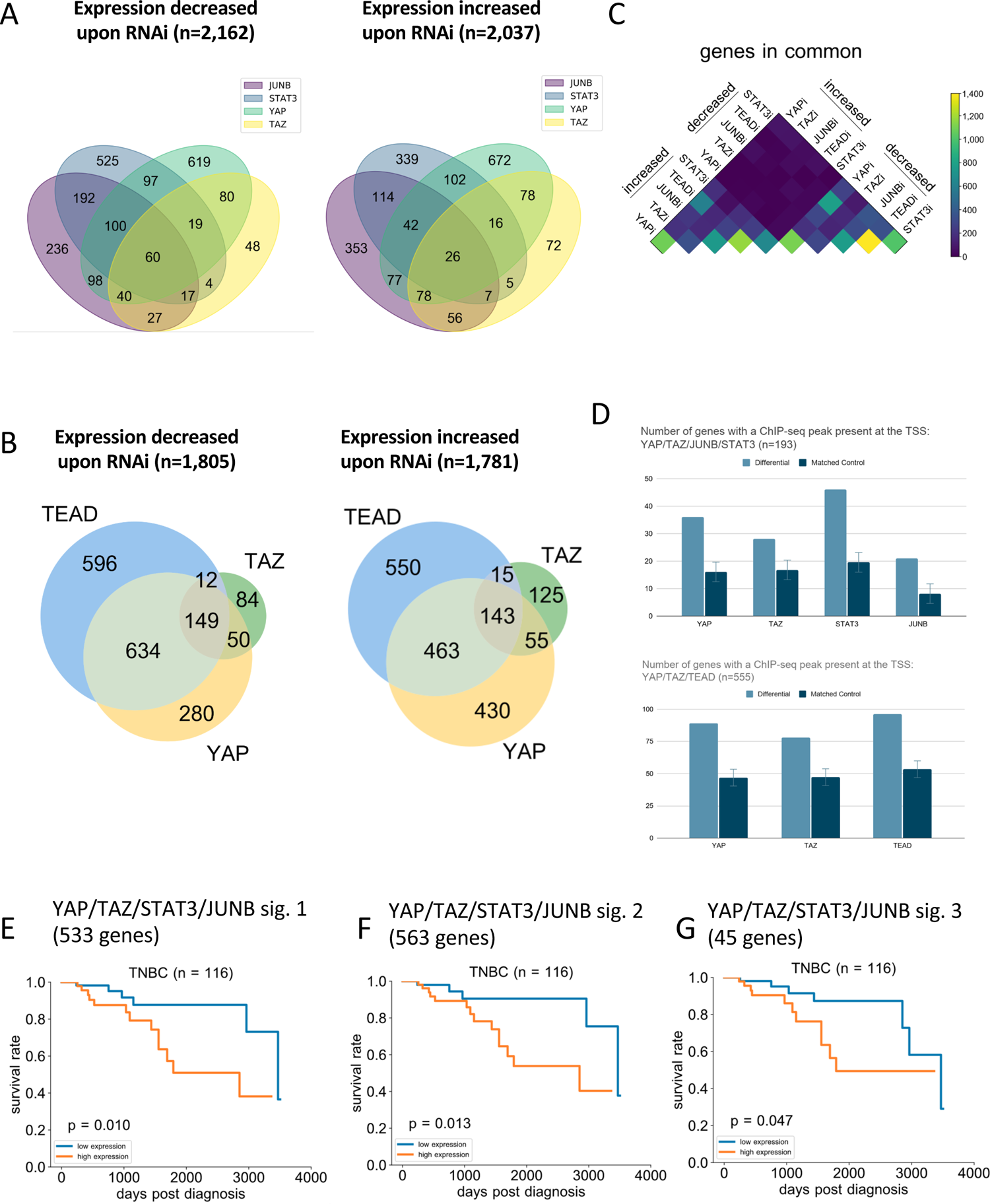
Core gene sets regulated by these factors are associated with differences in overall survival in triple-negative breast cancer patients. (A) Overlap between genes with decreased or increased expression in transformed cells treated with RNAi against YAP, TAZ, STAT3, or JUNB as compared with a control RNAi. (B) Same as (A), but comparing expression differences following RNAi against YAP, TAZ, and TEAD. (C) Heatmap showing overlap between genes upregulated and downregulated following RNAi; directionality of expression change is strongly conserved across RNAi conditions. (D) Number of differentially expressed (light blue) and matched random control (dark blue) genes having a binding site for the indicated transcription factors within 2 kb of at least one TSS. (E-G) Kaplan-Meier survival curves for three YAP/TAZ/STAT3/JUNB gene signatures in patients with high (orange) or low (blue) signature scores.

### YAP/TAZ gene signatures are associated with poor prognosis in triple-negative breast cancers

To determine the clinical significance of YAP/TAZ target genes in breast cancer samples, we first defined gene signatures impacted by YAP/TAZ/STAT3/JUNB or YAP/TAZ/TEAD (Table S4). For the YAP/TAZ/STAT3/JUNB set, we defined three gene signatures using the following criteria. The first set are target genes of YAP, TAZ, and JUNB in transformed condition that are also target genes of STAT3 in the transformed condition but not the non-transformed condition (i.e. near a tamoxifen-induced STAT3 peak). The second set are target genes of YAP, TAZ, and JUNB in the tamoxifen-treated condition that also have an AP-1 motif. The third set consists of genes from set 2 that are differentially expressed upon depletion of JUNB and either YAP or TAZ. For the YAP/TAZ/TEAD set, we defined the equivalents of sets 2 and 3 above, using TEAD peaks and motifs rather than JUNB peaks and AP-1 motifs.

The TCGA Invasive Breast Carcinoma Dataset includes 1,108 breast cancer samples from 1,101 patients with follow-up information, of which 116 are triple-negative breast cancers. For each signature, we computed a gene signature score (GSS) for each patient and then divided patients into low (GSS <0) and high (GSS > 0) gene signature expression (low and high risk). We evaluated the clinical significance of each gene signature by performing Kaplan-Meier analysis on overall survival, both on the entire set of patients and on subsets of patients with luminal A, luminal B, HER2+, and triple-negative breast cancer (TNBC) types.

The higher GSS groups for all three YAP/TAZ/STAT3/JUNB gene signatures are associated with a significantly shorter overall survival than the lower GSS group in TNBC patients (*p* value = 0.01, 0.01, and 0.04; Figure 7E-G). In contrast, survival was not significantly different for any of the three signatures for luminal A, luminal B, or HER2+ patients (Figure 7-figure supplement 1A-C). In addition, there is no significant difference in overall survival for the two YAP/TAZ/TEAD sets for any subset of cancer patients (Figure 7-figure supplement 2). As a control, we randomly selected a matched number of genes for each gene signature, and none these random gene signatures show significant differences in overall survival for any breast cancer subtype or for the entire cohort (Figure 7-figure supplement 1D-F show TNBC). These findings suggest the targeted gene signatures of YAP/TAZ and STAT3/JUNB complex are associated with poor prognosis in TNBC patients, which may have potential clinical application.

## DISCUSSION

### YAP and TAZ are transcriptional co-activators of AP-1 proteins and STAT3

YAP and TAZ, the major effectors of Hippo signal transduction pathway (Piccolo et al., 2014; Totaro et al., 2018; Ma et al., 2019), play critical roles in cancer initiation, progression, metastasis, and chemo-resistance in cancer therapy (Johnson and Halder, 2014; Yu et al., 2015; Zanconato et al., 2016). YAP/TAZ are transcriptional co-activators that are recruited by the TEAD family of transcription factors (Piccolo et al., 2014; Totaro et al., 2018; Ma et al., 2019; Moya and Halder, 2019). Here, we show that YAP and TAZ are important for transformation in ER-Src cells, which is mediated by an epigenetic switch involving an inflammatory regulatory network controlled by the joint action of NF-κB, STAT3, and AP-1 transcription factors (Iliopoulos et al., 2009; Iliopoulos et al., 2010; Ji et al., 2018; Ji et al., 2019). In this cellular model, YAP and TAZ co-associate with TEAD proteins at many target sites.

Here, we provide multiple lines of evidence demonstrating that YAP and TAZ are also transcriptional co-activators of STAT3 and AP-1 proteins. First, YAP and TAZ co-immunoprecipitate with JUNB and STAT3 in nuclear extracts. Second, all direct pairwise interactions between YAP or TAZ with JUNB or STAT3 are observed with proteins expressed in *E. coli*. The WW1 domain of YAP and the WW domain of TAZ are critical for these interactions and for transformation, but they are dispensable for the interaction with TEAD proteins. Third, YAP and TAZ co-associate with JUNB and/or STAT3 at many target sites with peak summits very close together. Similar results are observed in a triple-negative breast cell line (MDA-MB-231). Fourth, YAP/TAZ target sites coincide with AP-1 and, to a lesser extent, STAT3 motifs. Indeed, the frequency of AP-1 motifs among YAP/TAZ target sites is comparable to that of TEAD motifs. Fifth, sequential ChIP experiments directly indicate that YAP/TAZ co-occupy target sites with JUNB and STAT3.

### Specificity of YAP and TAZ recruitment to target sites

Although YAP and TAZ directly interact with STAT3 and JUNB, the specificity of YAP/TAZ association with genomic sites is complex and poorly understood. YAP/TAZ are only recruited to a subset of genomic regions bound by STAT3 and/or AP-1, indicating that the direct physical interactions are not sufficient for recruitment. With respect to the target sites and hence the DNA-bound proteins, YAP/TAZ associate with AP-1 and TEAD motifs at comparable frequency, but less frequently with STAT3 motifs. In addition, STAT3 and AP-1 proteins directly interact, co-associate with many genomic regions through AP-1 motifs and affect the expression of common genes (Ji et al., 2019). Thus, it is difficult to determine the contributions of AP-1 proteins and STAT3 to YAP/TAZ recruitment, although these DNA-binding activator proteins play a role individually and in combination. On the other hand, NF-κB is not involved in YAP/TAZ recruitment, even though it can act together with STAT3 and AP-1 factors to mediate the inflammatory regulatory network involved in transformation (Ji et al., 2019). We presume that, in addition to AP-1 proteins and STAT3, YAP/TAZ recruitment to specific genomic regions is influenced by other proteins and/or DNA sequences near the AP-1 and/or STAT3 sequence motifs.

YAP/TAZ, via its interaction with TEAD proteins, can synergize with AP-1 proteins at composite regulatory elements containing both TEAD and AP-1 motifs (Zanconato et al., 2015). In this regard, we observe a small subset of YAP/TAZ target sites that contain both AP-1 and TEAD motifs. However, most YAP/TAZ target sites that contain AP-1 or STAT3 motifs do not contain a TEAD motif and vice versa. Thus, YAP/TAZ can act as a co-activator for AP-1 and STAT3 in a manner that is independent of TEAD proteins.

Although YAP and TAZ have extremely similar binding profiles, they are not identical. In particular, we used two independent analyzes to identify 450 TAZ-specific target sites and 125 YAP-specific target sites. Interestingly, the TAZ-specific sites are enriched for ETS motifs and depleted for AP-1 motifs; TEAD motifs neither enriched or depleted. These observations suggest an ETS protein(s) can interact specifically (although not necessarily directly) with a region of TAZ that is not found in YAP. In addition, the TAZ-specific sites are enriched as certain classes of genes, suggesting a distinct biological function.

Lastly, about half of the YAP/TAZ target sites are not associated with AP-1, STAT3, or TEAD motifs suggesting that other transcription factors can recruit YAP/TAZ. CEBP motifs are enriched and frequently observed at YAP/TAZ target sites in ER-Src cells, but not in MDA-MB-231 cells. Conversely, RUNX3 motifs are enriched at YAP/TAZ target sites in MDA-MB-231 cells, but not in ER-Src cells. Such cell-line specificity could be due to differences in the amount or activity of the recruiting protein(s) that recognize the motif and/or differences in auxiliary proteins that are important for recruitment to these motifs.

### YAP/TAZ targets associated with AP-1 proteins and STAT3 have different functions and cancer properties from those associated with TEAD proteins

The current view is that the biological functions of YAP/TAZ are mediated primarily through their interaction and recruitment by TEAD proteins (Piccolo et al., 2014; Totaro et al., 2018; Ma et al., 2019; Moya and Halder, 2019). However, our results indicate that a comparable number of YAP/TAZ targets involve recruitment by AP-1 proteins and/or STAT3. The gene signatures of these two classes of YAP/TAZ targets are different. TEAD-associated sites are enriched for genes associated with cell motility and cytoskeletal organization, but they are not associated with a significant difference in overall survival for breast cancer patients. In contrast, the gene signature of AP-1/STAT3-associated YAP/TAZ targets is not associated with cell motility genes but is linked to shorter survival of triple-negative breast cancer patients; no overall survival difference is observed for other breast cancer subtypes. Thus, in addition to serving as a coactivator for TEAD proteins, YAP and TAZ are also co-activators for AP-1 proteins and STAT3, with both classes of target sites and affected genes playing important, yet distinct, roles in cancer-related phenotypes and cancer progression.

## MATERIALS AND METHODS

### Cell lines and chemicals

MCF-10A-ER-Src cells (Iliopoulos et al., 2009; Iliopoulos et al., 2010) were grown in DMEM/F12 without phenol red (Thermo Fisher Scientific, 11039-047) + 5% charcoal stripped FBS (Sigma, F6765) + 1% pen/strep (Thermo Fisher Scientific, 15140122)+20 μg/ml Hydrocortisone (Sigma, H-0888) + 0.1 μg/ml insulin (Sigma, 10516). 1-0.4 μM AZD0530 (Selleck Chemicals, S1006) were used to μ induce the transformation. MDA-MB-231 cells were grown in DMEM (Thermo Fisher Scientific, 11995-073) + 10% FBS (Sigma, TMS-013-B) + 1% pen/strep.

### Cell transformation assays

The transformation capacity was measured by growth in low attachment conditions (GILA) (Rotem et al., 2015) or in soft agar. For the soft agar assay, 10^4^ cells in culture medium were mixed with 0.4% low melting point agarose (VWR, 89125-532) at 37^0^C and seeded on top of 1% agarose standing layer in 12 well dishes. Colony density was measured 2-3 weeks after seeding with images captured by a digital camera (Olympus SP-350; Cam2com). For the GILA assay, 2000 cells were seeded into ultra-low attachment surface 96-well plate (Costar, 3474). 5 days after seeding, sphere cells growing in the low attachment plates were quantitated by CellTiter-Glo luminescent cell viability assay (Promega, G7571) using a SpectraMax M5 Multi-Mode Microplate Reader (Molecular Devices). Cells growing in regular culture dishes were stained using crystal violet and cell density was measured using Fiji Image J’s (version 1.52b) measurement function.

### CRISPR knockout and siRNA knockdown

CRISPR sgRNAs, designed with previously described algorithms (Hsu et al., 2013), were cloned into a CRISPR-blasticidin lentiviral plasmid, which was constructed by replacing puromycin resistant gene with blasticidin resistant gene of LentiCRISPR V2 plasmid (Addgene, #52961). The oligo sequences used to clone into CRISPR vector are: YAP exon 8—AAACTCTCATCCACACTGTTCAGGC and CACCGCCTGAACAGTGTGGATGAGA; TAZ exon 3— AAACCCCGACGAGTCGGTGCTGGAC and CACCGTCCAGCACCGACTCGTCGGG.

CRISPR lentiviral plasmids and VSV-G, GP and REV plasmids were transfected into 293T cells to produce CRISPR lentivirus. CRISPR lentiviruses infected ER-Src cells for 1 day, and μg/ml blasticidin (Thermo Fisher, R21001) for additional 3 days.

Oligo siRNAs were purchased from Dharmacon (siGENOME SMART pool) (Table S5) and were transfected into ER-Src cells using Lipofectamine RNAiMAX (Thermo Fisher Scientific, 13778050). 24 hours after transfection, cells were split and then were treated with tamoxifen and AZD0530 (4 μM) for additional 24 hours before the following assays.

### Cell Fractionation, Co-IP and Western Blot

Cell fractionation protocols were similar to the protocols as described before with some changes (Wu et al., 2002). Cells were washed with cold PBS, resuspended using buffer A (10 mM HEPES pH 7.5, 10 mM KCl and 2 mM MgCl2), lysed by grinding 50 times in Wheaton A (Wheaton, 357538), and then incubated on ice for 20 minutes. The lysate was placed on top of 30 % sucrose and pelleted by spinning at 15000 RPM at 4°C for 10 minutes. After removing the supernatant as the cytoplasm fraction, pellets were resuspended in buffer GB (20 mM TrisCl pH7.9, 50% glycerol, 75 mM NaCl, 0.5 mM EDTA and 0.35 mM DTT) and then mixed with an equal amount of buffer NLB (20 mM HEPES pH7.6, 300 mM NaCl, 7.5 mM MgCl2, 1% NP40, 1 M Urea, 0.2 mM EDTA and 1 mM DTT) and incubated on ice for 2 min. After centrifugation at 15000 RPM for 5 minutes, supernatants representing the nucleoplasm fraction were removed, and pellets resuspended using protein lysis buffer (20 mM Tris pH 7.4, 150 mM NaCl, 1 mM EDTA, 1 mM EGTA, 1% Triton X-100, 25 mM sodium pyrophosphate, 1 mM NaF, 1 mM β μg/ml aprotinin). The material was sonicated 4×15 seconds using Branson Microtip Sonifier 450 at 60% cycle duty and 4.5 output and spun at 15000 RPM for 5 minutes to obtain the soluble chromatin fraction.

Co-immunoprecipitations were performed as described previously (Ji et al., 2019). Briefly, lyates were mixed with antibodies and 10 μl co-IP buffer (50 mM Tris PH 7.5, 100 mM NaCl, 1.5 mM μ EGTA and 0.1% Triton X-100), then rotated at 4 ^0^C overnight, and washed with a co-IP buffer 8 times. Antibodies used for co-IP and western blot can be found in the Table S5.

### qPCR

RNA was extracted using mRNeasy Mini Kit (Qiagen, No. 217004). 1 μ RNA was converted to cDNA using SuperScript III Reverse Transcriptase (Thermo Fisher Scientific, 18080093). qPCR was running using a 7500 Fast Real-time PCR system (Applied Biosystems). qPCR primer sequences can be found in the Table S5. All experiments were run independently three times and values of each mRNA were normalized to that of the 36b4 internal control gene.

### ELISA assay for IL-6 secretion

IL-6 secretion was measured using human IL-6 immunoassay kit (R&D Systems, D6050) as per manufacturer’s instructions. Optical density was determined using SpectraMax M5 Multi-Mode Microplate Reader (Molecular Devices).

### Luciferase reporter assay

The pGL-AP-1 plasmid containing 6 consensus AP-1 binding sites was co-transfected with the pRL-CMV plasmid (Promega) into cells using TransIT 2020 transfection reagent (Mirus Bio, MIR5400). A pRL-CMV plasmid expressed Renilla protein and was used as an internal transfection control. 24 hours after transfection, cells were split and treated with tamoxifen for 24 h to induce transformation. After 3 days, firefly and Renilla luciferase activities were determined by Dual-luciferase Reporter Assay kit (Promega, E1910). To evaluate NF-κB activity, pGL4.32 [luc2P/NF-κB-RE/Hygro] (E8491, Promega) and pRL-CMV were co-transfected into cells and the same protocol followed.

### Recombinant proteins and direct interactions by co-IP

To produce recombinant proteins, YAP, TAZ, STAT3 and JUNB were cloned into pET30 plasmid. Recombinant proteins were produced in BL21 *E. Coli* and dialyzed in a neutral buffer (150 mM NaCl, 46.6 mM Na2HPO4 and NaH2PO4 pH 8.0). The direct interactions among these recombinant proteins were examined *in vitro* using a similar co-IP procedure as mentioned above.

### ChIP-seq

Cells were dual cross-linking with a mixture of 2 mM each of ethylene glycol bis (succinimidyl succinate) (EGS) and disuccinimidyl glutarate (DSG) and 1% formaldehyde. Chromatin was digested with 60 units MNase (New England Biolabs, M0247S) at 37 ^0^C for 10 minutes and then sonicated using Branson Microtip Sonifier 450 (4X15 second at output 4.5 and duty cycle 60%) to the sizes mostly between 150-500 bp. 50 μ antibodies for transcription factors (listed in Table S5), and 15 μ (Thermo Fisher Scientific, 10004D) was used for the chromatin immunoprecipitation (ChIP). ChIP-seq libraries were sequenced using Hiseq 2000 at the Bauer Core Facility, Harvard.

### Sequential ChIP

Sequential ChIP was performed and analyzed as described previously (Geisberg and Struhl, 2004; Miotto and Struhl, 2011). 100 μg of chromatin was mixed with 20 μl Dynabead and μl ChIP IP buffer (20 mM TrisCl pH 7.5, 140 mM NaCl, 2 mM EDTA and 2 mM EGTA), rotated at 4 ^0^C overnight, and sequentially washed with 1 ml each of ChIP IP buffer, ChIP wash buffer I (20 mM TrisCl pH 7.5, 140 mM NaCl, 2 mM EDTA, 2 mM EGTA and 0.5% Triton-100), ChIP wash buffer III (twice, 20 mM TrisCl pH 7.5, 250 mM LiCl, 2 mM EDTA, 2 mM EGTA and 0.5% Triton-100) and TE μl sequential ChIP elution buffer (20 mM TrisCl pH 7.5, 500 mM NaCl, 2 mM EDTA, 2 mM EGTA 30 mM DTT, 0.1% SDS and cOmplete protease inhibitor cocktail (Roche, 11873580001) at 37 ^0^C for 30 minutes. Eluted chromatin l Dynabeads and antibodies in 200 μ ChIP IP buffer and then rotated at 4 ^0^C overnight. Chromatin from the second ChIP were washed, eluted, and de-crosslinked. Chromatin obtained from single ChIP and sequential ChIP samples were analyzed using qPCR. ChIP primers are listed in Table S5.

### ChIP-seq analysis

FASTQ reads were aligned to the human reference genome (GRCh38) using Bowtie2 (Langmead and Salzberg, 2012). For post-alignment filter steps, we used samtools-1.9 with MAPQ threshold 30 and picard-tools-2.18 to remove low quality and duplicate reads. The SPP algorithm with --cap-num-peak 300000 and IDR-2.0.4 (Landt et al., 2012) with --soft-idr-threshold 0.05 was then used to determine the genomic binding sites of each transcription factor. The number of IDR peaks are listed in Table S1. Peaks within 2 kb of a GENCODE TSS are denoted as TSS-proximal; others are denoted as distal. DESeq2 (Love et al., 2014) was used to define protein-specific peaks between YAP and TAZ with the cutoff log_2_ fold change > 0 and multiple-testing adjusted *p*-value < 0.05.

Occupancy analysis we performed using IDR peaks. We used the MACS2 pileup command to generate signal profiles which were smoothed using a 10 base pair Gaussian kernel to reduce noise. We then identified summits as the positions within a peak with the highest signal value. We adjusted IDR peaks to the summits ± 150 regions to avoid the potential bias due to varied ChIP-seq peak length distribution between transcription factors. Bedtools-2.29.2 was used to identify overlapped regions and calculate distance between peak summits.

We downloaded DNase hypersensitivity sites from the ENCODE Portal (ENCODE, 2020) for ER-Src cells treated for 24 h with tamoxifen (ENCODE accession ENCSR752EPH) and used bedtools intersect to determine that YAP/TAZ are present at 47,547 MCF10A DHSs. To assess co-occupancy of factors with YAP/TAZ for statistical significance, we compared the observed fractions of peak intersection with these YAP/TAZ-bound DHSs against a control set of 47,547 DNase hypersensitivity sites in ER-Src cells that are not bound by YAP/TAZ. We generated the control set as follows: (1) we obtained DNase-seq signal Z-scores for 2 million human DHSs from the ENCODE Encyclopedia (ENCODE, 2020) in tamoxifen-treated MCF10A cells (ENCODE accession ENCSR752EPH); (2) we identified DHSs from this set intersecting YAP/TAZ IDR peaks; (3) we selected 47,547 random DHSs from the ENCODE collection with the same Z-score distribution as the DHSs which intersect YAP/TAZ IDR peaks; (4) we iteratively replaced any randomly selected DHSs which intersected YAP/TAZ peaks with alternative DHSs of the same Z-score until none of the 47,547 intersected YAP/TAZ peaks.

For motif occurrence analysis, TEAD, JUNB, STAT3, CEBPA, NF-κb motifs were downloaded from the HOCOMOCO database (Kulakovskiy et al., 2018). We used FIMO (Bailey et al., 2015) to scan the human genome for motif matches with a specific threshold for each motif as a function of the information content of the motif. For motif enrichment analysis, we downloaded the complete set of transcription factor motifs in the HOCOMOCO database (Kulakovskiy et al., 2018) and used FIMO to scan IDR peaks for each transcription factor (Machanick and Bailey, 2011). We then generated control DHS sets for each set of IDR peaks as described above, scanned these sets for motif occurrences using FIMO, and computed enrichment Chi-square *p*-values for each motif. We used this same workflow for TAZ-specific peaks.

GO analysis was performed using GOrilla (Eden et al., 2009). For YAP/TAZ-specific peaks, two different control gene sets were tested: (1) genes proximal to non-specific YAP/TAZ peaks and (2) genes proximal to any active DHS in MCF10A cells. The results were similar; results from the former approach are reported. For YAP/TAZ peaks separated by motif, the complete set of genes proximal to YAP/TAZ peaks was used as the control set. For non-specific YAP/TAZ peaks, the complete set of genes proximal to any active DHS in MCF10A cells was used as the control set.

### RNA-seq analysis

RNA was prepared used mRNeasy Mini Kit (Qiagen, No. 217004). 0.4 μg total RNA was used for RNA-seq library preparation. RNA-seq libraries were generated using TruSeq Ribo Profile Mammalian Kit (Illumina, RPHMR12126). RNA-seq libraries were sequenced at Bauer Core Facility using Hiseq 2000. For analysis, we trimmed adapter sequences, ambiguous ‘N’ nucleotides (the ratio of “N” > 5%), and low-quality tags (quality score < 20). Clean reads were aligned against the GENCODE v30 reference transcriptome (Harrow et al., 2012) using STAR (Dobin et al., 2013) with the following parameters:

--outFilterMultimapNmax 20

--alignSJoverhangMin 8

--alignSJDBoverhangMin 1

--outFilterMismatchNmax 999

--outFilterMismatchNoverReadLmax 0.04

--alignIntronMin 20

--alignIntronMax 1000000

--alignMatesGapMax 1000000

--sjdbScore 1

Gene counts were normalized to TPM (transcripts per million RNA molecules) using RSEM (Li and Dewey, 2011) with the following parameters: “--estimate-rspd --calc-ci.” Differential expression analysis was performed with DESeq2 (Love et al., 2014) with default parameters. A separate DESeq2 run was performed for siRNA against each factor, comparing replicates treated with the siRNA against the given factor versus replicates treated with the control siRNA. DESeq2 was performed independently for tamoxifen-treated MCF10A and ethanol-treated MCF10A for each siRNA. Genes with a multiple-testing adjusted *p*-value <0.01 were considered to be differentially expressed.

To assess whether differential genes were significantly more likely to be directly regulated by the transcription factors of interest, we divided differential genes into two classes: TF target genes, having an IDR peak for a given factor within 2 kb of any of the gene’s transcription start sites based on GENCODE v30 annotations; and non-TF target genes, not having a peak within that distance of any TSS. We compared each set of differential genes against a control set of the same number of protein coding genes randomly selected from the GENCODE annotations. A chi-square *p*-value was computed for the respective fractions of target and non-target genes.

### Gene signature and survival analysis in breast cancer samples from TCGA

TCGA Provisional data, including 1,108 breast cancer samples, were obtained from https://www.cbioportal.org. These samples were sub-divided into Luminal A, Luminal B, HER2+, and TNBC groups according to ER/PR/HER2 status based on immunohistochemistry (Parker et al., 2009; Wallden et al., 2015). We used the lifelines Python package to perform Kaplan-Meier survival analysis. Gene signature score (GSS) were calculated using this formula: GSS=∑(x_i_-μ_i_)/σ_i_, where x_i_ is the expression of i gene in patient samples; μ_i_ is mean of i gene in all patient samples; σ_i_ is standard deviation. Low expression (low risk) group was GSS<0 and high expression (high risk) group was GSS ^≥^0 (Adorno et al., 2009). We refined the gene signatures by computing the coefficient of variation (CV) of the expression of all genes among these patients, removing genes with little (CV < 5%) or excessive variation (CV > 85%), the latter being likely to be statistical noise (Mar et al., 2011; Jang et al., 2019), and then filtering out pseudogenes (Table S4).

In defining gene signatures, we considered an individual gene to be targeted by a transcription factor if a ChIP-seq peak for that factor fell within 2 kb of at least one of its transcription start sites from the GENCODE catalog (version 30). We considered an individual gene to be differentially expressed following RNAi treatment against a given factor if its multiple-testing adjusted *p*-value from DESeq2 was less than 0.01.

Three YAP/TAZ/STAT3/JUNB gene signatures were defined as follows. The first signature was defined by (1) identifying all targeted by all three of YAP, TAZ, and JUNB in tamoxifen-treated MCF10A cells using the above criteria for ChIP-seq, then (2) filtering these to contain only genes which are targeted by STAT3 in TAM-treated MCF10A cells but not ETH-treated MCF10A cells based on ChIP-seq. The second signature was defined by (1) identifying all genes targeted by all three of YAP, TAZ, and JUNB in tamoxifen-treated MCF10A cells using the above criteria for ChIP-seq, then (2) filtering these to contain only genes for which the associated YAP/TAZ peaks contain a JUNB motif. The third signature was defined by filtering genes from the second signature to require that they be differentially expressed following JUNBi treatment and also differentially expressed following either YAPi treatment, TAZi treatment, or both.

Three YAP/TAZ/TEAD gene signatures were defined as follows. The first signature was defined by identifying all genes targeted by all three of YAP, TAZ, and TEAD in tamoxifen-treated MCF10A cells using the above criteria for ChIP-seq. The second signature was defined by filtering genes from the first signature to contain only genes for which the associated YAP/TAZ peaks contain a TEAD motif. The third signature was defined by filtering genes from the second signature to contain only genes differentially expressed following TEADi treatment and also differentially expressed following either YAPi treatment, TAZi treatment, or both.

### Data deposition

All sequencing data were deposited on National Cancer for Biotechnology Information Gene Expression Omnibus (GEO). GSE166943 is the accession number for all the data, with GSE166941 being the subset for the ChIP-seq data and GSE166942 for the RNA-seq data.

## Supporting information

All supplemental figures

## ACKNOWLEDGEMENTS

We thank Zhe Ji and Xueli S. Wu for help in the initial stages of bioinformatic analysis. This work was supported by grants from the National Institutes of Health to Z.W (HG009446) and K.S. (CA 107486).

## SUPPLEMENTARY FIGURE LEGENDS

**Figure 2-figure supplement 1.** Gene expression in siRNA knockdowns. RNA levels of YAP, TAZ, STAT3, and NF-κB1 in non-transformed (ETH) or transformed (TAM) cells knocked-down by siRNA (i) of the indicated factors.

**Figure 3-figure supplement 1.** Interactions of YAP and TAZ with JUNB and STAT3. (A) Levels of the indicated proteins in the cytoplasm or nucleus of non-transformed (E) or transformed (T) cells. LDH-A is a cytoplasm marker and H3 is a nucleus marker. (B) Levels of the indicated proteins upon immunoprecipitation (IP) with antibodies (Anti) against the indicated proteins and the IgG control. The Anti-YAP/TAZ antibody recognizes both YAP and TAZ. (C) Growth of cells overexpressing the indicated proteins (EV is the empty vector control) under conditions of high attachment; n = 3 with error bars: ±SD.

**Figure 4-figure supplement 1.** Motif enrichment and overlap of ChIP-seq peaks. Enrichment of the indicated motifs (different colors) in STAT3, JUNB, TEAD, and TAZ-specific (i.e. not bound by YAP) binding sites as compared with matched control DNase-hypersensitive regions in transformed cells.

**Figure 5-figure supplement 1.** Co-binding of YAP/TAZ with other factors. (A) Intersection of various combinations of binding sites for the four factors in transformed cells. The number of sites for each factor is indicated and black horizontal bars at the bottom left of the panel. (B) Distance between peak summits for biological replicates or combinations of the indicated factors in non-transformed cells. (C) Sequential ChIP of YAP or TAZ with JUNB in the indicated orders in untransformed (ETH) and transformed (TAM) cells at the SNX24 and MYC loci. (D) Sequential ChIP of YAP or TAZ with JUNB in the indicated orders in untransformed (ETH) and transformed (TAM) cells at the MYC locus.

**Figure 6-figure supplement 1.** Venn diagrams of the overlap of the indicated transcription factor binding sites in non-transformed (ETH) and transformed (TAM) ER-Src cells and in MDA-MB-231 cells.

**Figure 6-figure supplement 2.** YAP/TAZ binding sites and their co-binding with other factors in MDA-MD-231 cells. (A) Correlation between YAP and TAZ binding levels, with YAP-specific (green) and TAZ-specific (yellow) sites highlighted. (B) Intersection of various combinations of binding sites for the four factors in ethanol-treated MDA-MB-231 cells. (C) Inter-summit distances for biological replicates and pairwise combinations of binding sites in MDA-MB-231 cells.

**Figure 6-figure supplement 3.** YAP/TAZ Motif enrichment and co-binding of YAP/TAZ with other factors in MDA-MD-231 cells. (A) Percent of binding regions for the indicated proteins that contain a given DNA sequence motif (different colors). (B) Enrichment of motifs within YAP/TAZ binding sites as compared with control DHSs. (C) Histogram of all motif locations for TEAD, JUNB, STAT3, NF-κB, and CEBPB occurring within 500 bp of a YAP (top) or TAZ (bottom) peak summit (defined as position 0). (D) Intersection between YAP and TAZ sites containing STAT3, AP-1, and TEAD motifs. (E) Enrichment of motifs within TAZ-specific peaks as compared with control DHSs.

**Figure 6-figure supplement 4.** Genes targeted by YAP/TAZ and associated factors based on ChIP-seq data. (A) Target genes of YAP/TAZ as determined by proximity of a TSS to a ChIP-seq peak in tamoxifen-treated ER-Src or MDA-MB-231 cells. (B) Overlap between JUNB, STAT3, YAP, and TAZ binding sites in target genes in transformed ER-Src cells (C) Overlap between YAP/TAZ target genes having a CEBP, STAT, JUN, or TEAD motif proximal to at least one TSS in transformed ER-Src cells. (D) GO enrichment terms for genes targeted by a YAP/TAZ peak containing a CEBP motif (top) or a TEAD motif (bottom) in tamoxifen-treated MCF10A cells.

**Figure 7-figure supplement 1.** Kaplan-Meier survival curves for (A) YAP/TAZ/STAT3/JUNB gene signature 1, (B) YAP/TAZ/STAT3/JUNB gene signature 2, and (C) YAP/TAZ/STAT3/JUNB gene signature 3 for luminal A (left), luminal B (center), or HER2+ (right) breast cancer. (D-F) Kaplan-Meier survival curves for a random control set of genes numbering the same as the three YAP/TAZ/STAT3/JUNB gene signatures in triple-negative breast cancer.

**Figure 7-figure supplement 2.** Kaplan-Meier survival curves for (A) YAP/TAZ/TEAD gene signature 1, (B) YAP/TAZ/TEAD gene signature 2 for luminal A, luminal B, HER2+, and triple-negative breast cancer patient cohorts.

## SUPPLEMENTARY TABLE LEGENDS

**Table S1.** Correlation between (A) biological replicates and (B) number of binding sites of the indicated proteins in ER-Src (ethanol or tamoxifen; wild-type and YAP or TAZ knockout (KO) and MDA-MB-231 cells..

**Table S2.** Enriched GO categories for YAP-specific, TAZ-specific, or shared YAP/TAZ target genes.

**Table S3.** Enriched GO categories for YAP/TAZ target sites containing the indicated motifs.

**Table S4.** Gene signatures for the indicated categories of YAP/TAZ target sites.

**Table S5.** siRNAs, antibodies, qPCR primers, ChIP-seq primers, and list of datasets.

