## Supplementary material for "YAP and TAZ are transcriptional co-activators of AP-1 proteins and STAT3 during breast cellular transformation": All supplemental figures

**Figure 2-figure supplement 1**

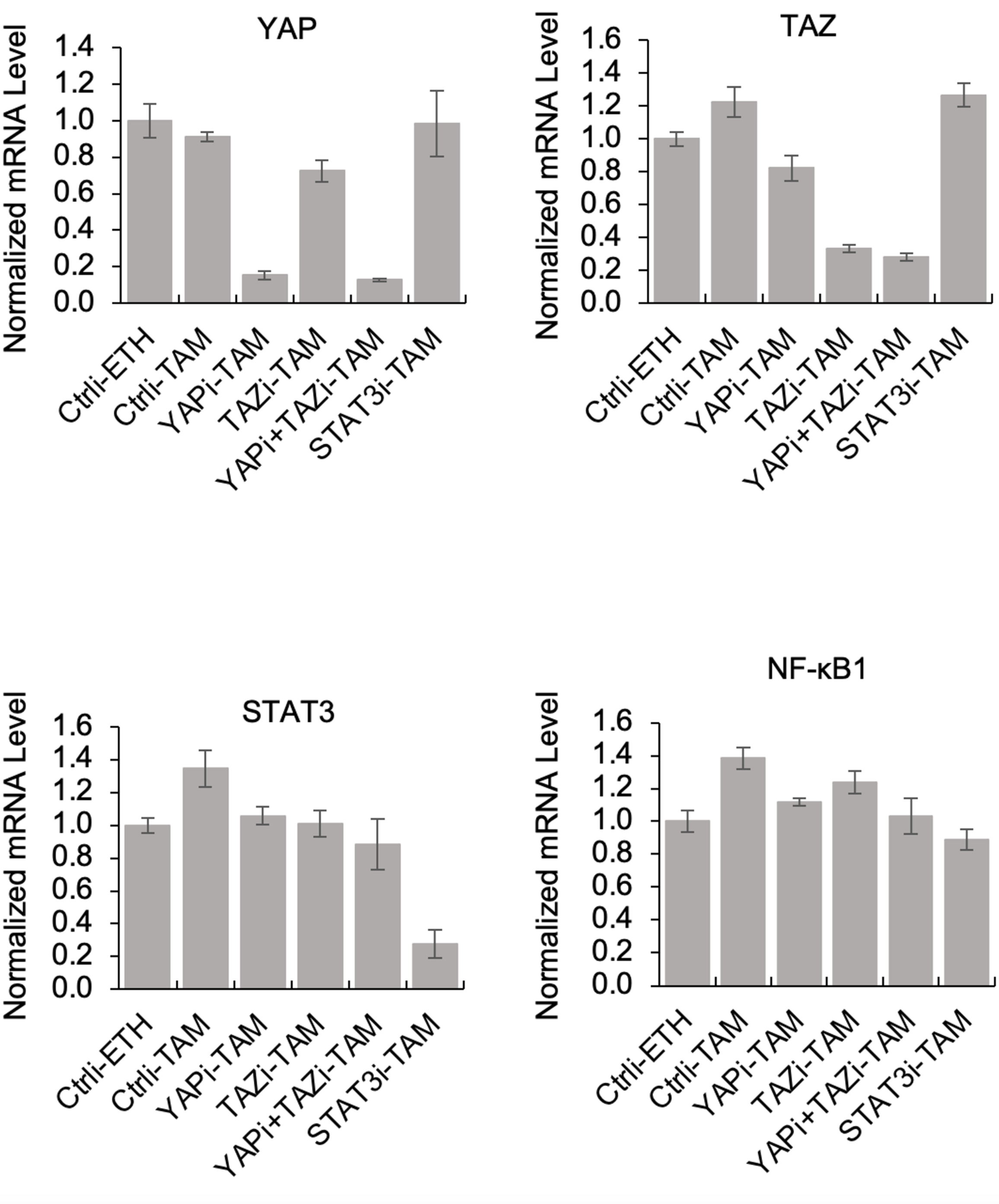

**Figure 3-figure supplement 1**

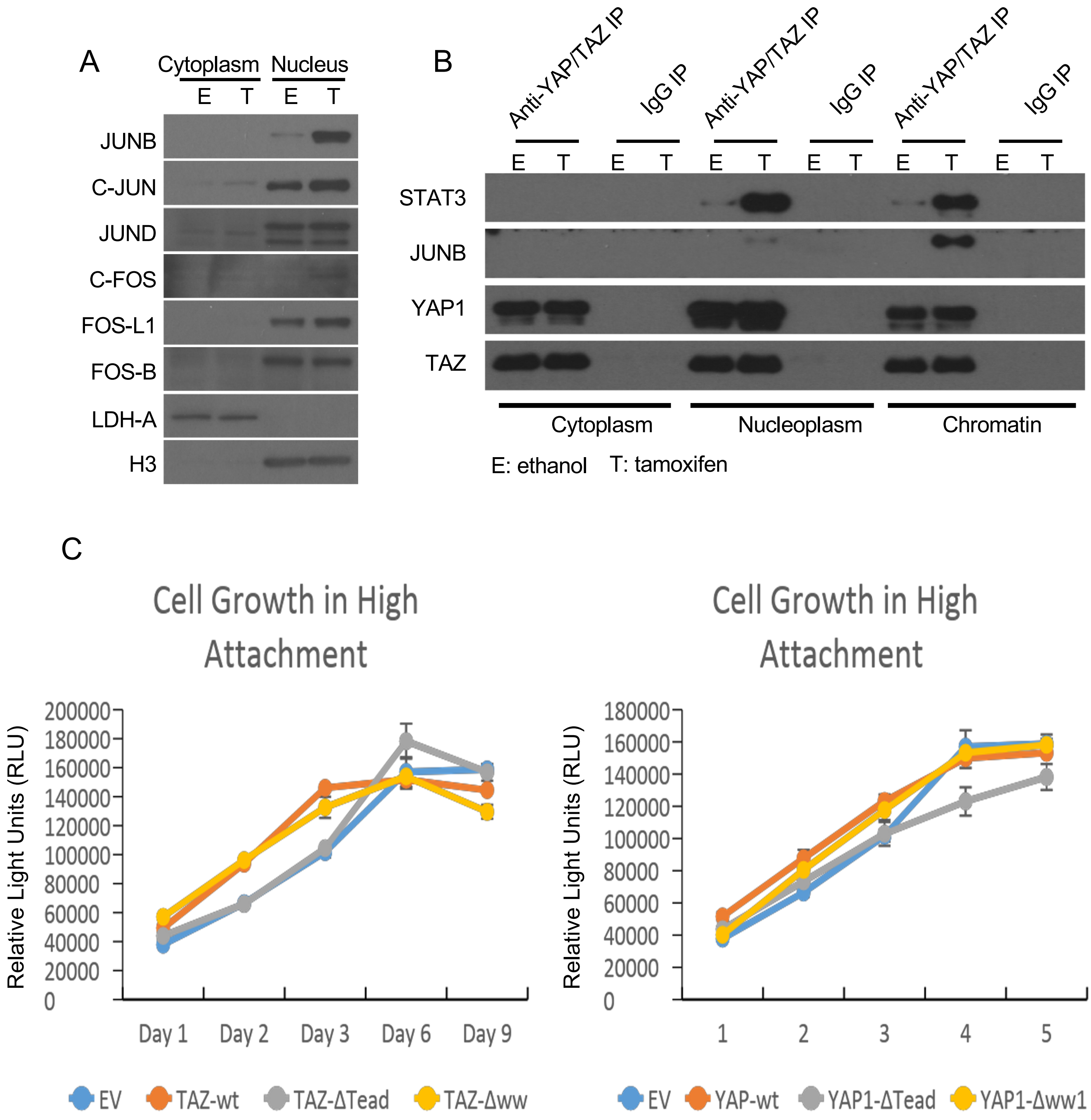

**Figure 4-figure supplement 1**

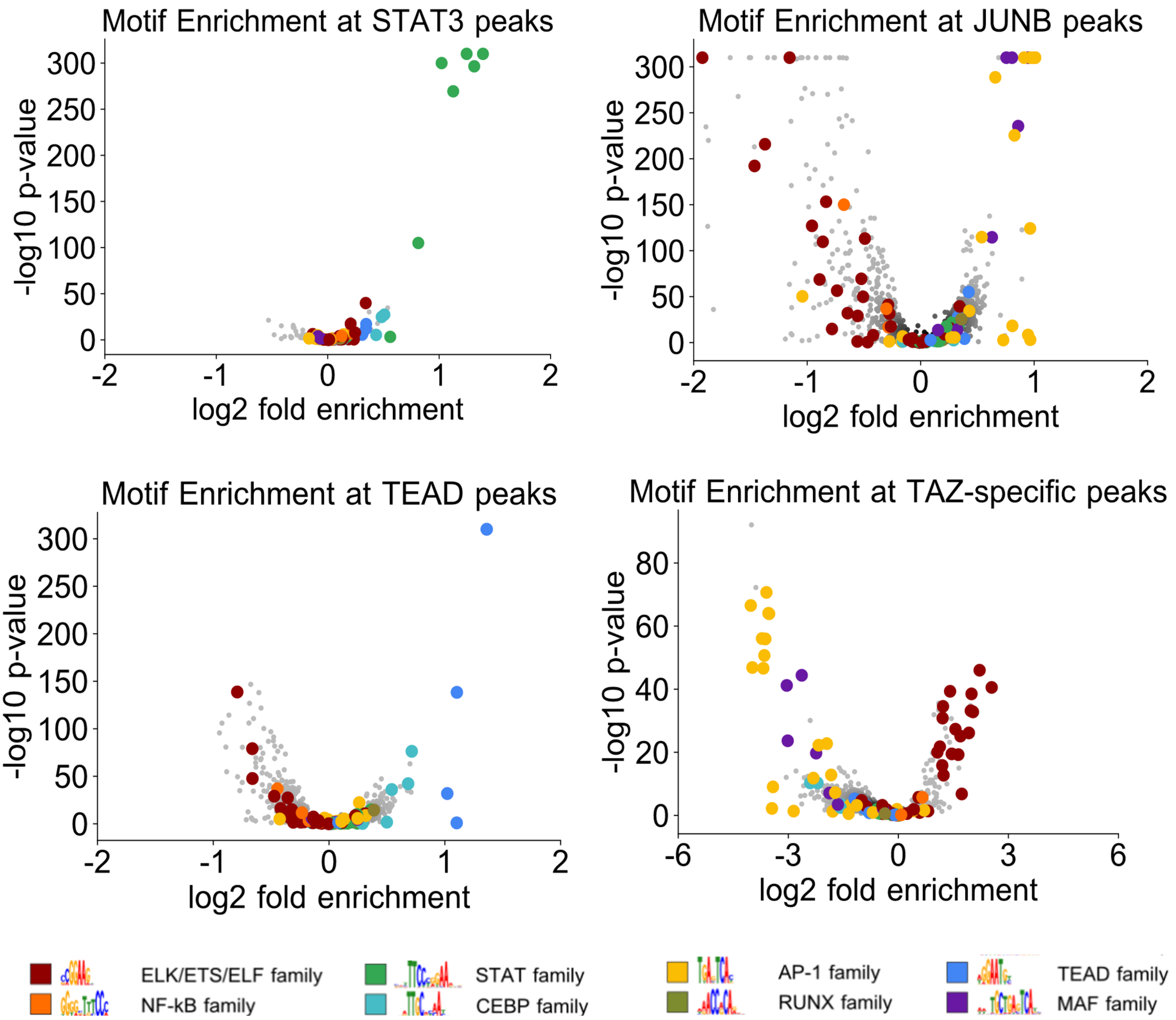

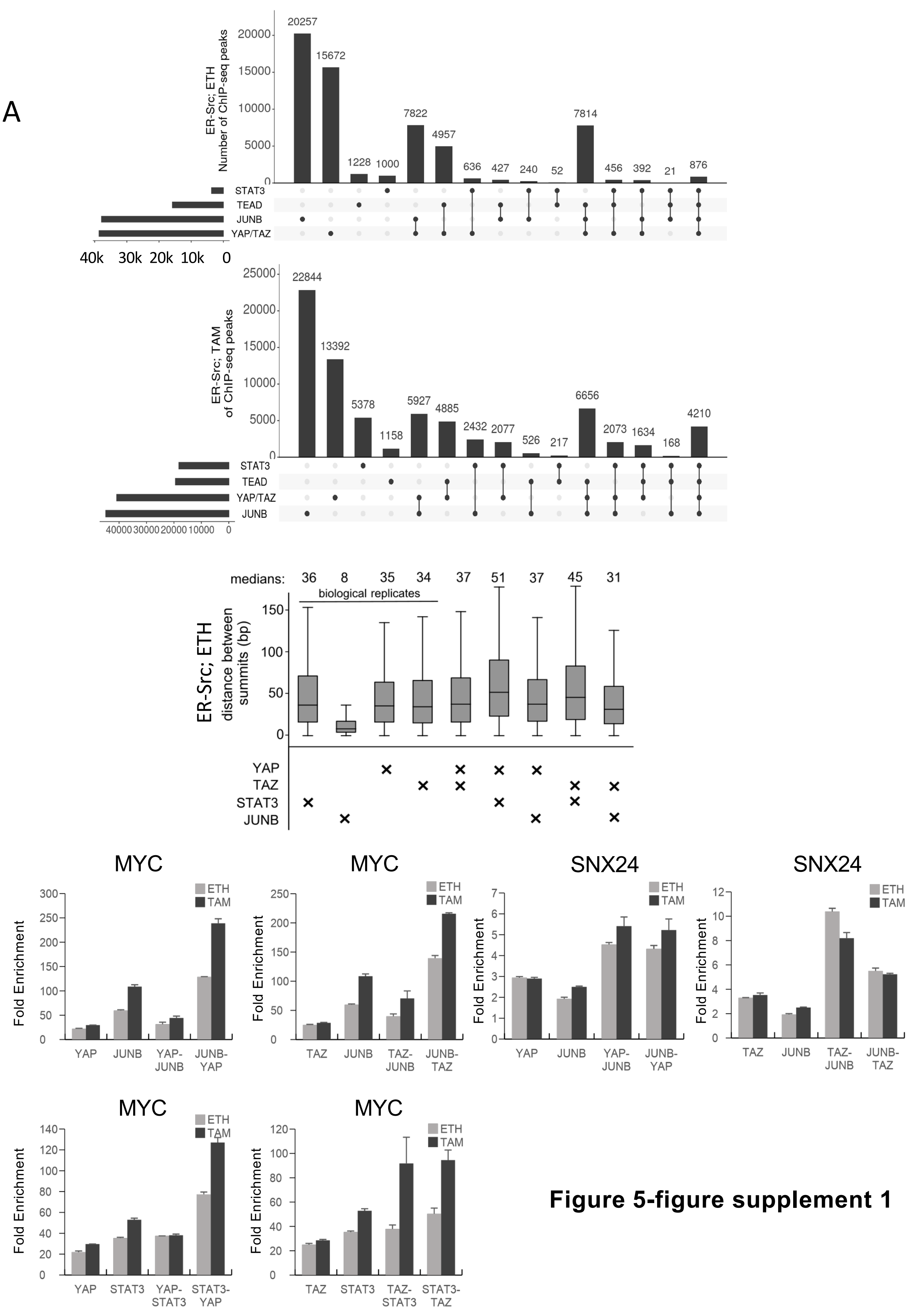


**Figure 5-figure supplement 1**

#### Figure 6-figure supplement 1

**Overlap of ChIP-seq Peaks in Three Cell Lines**

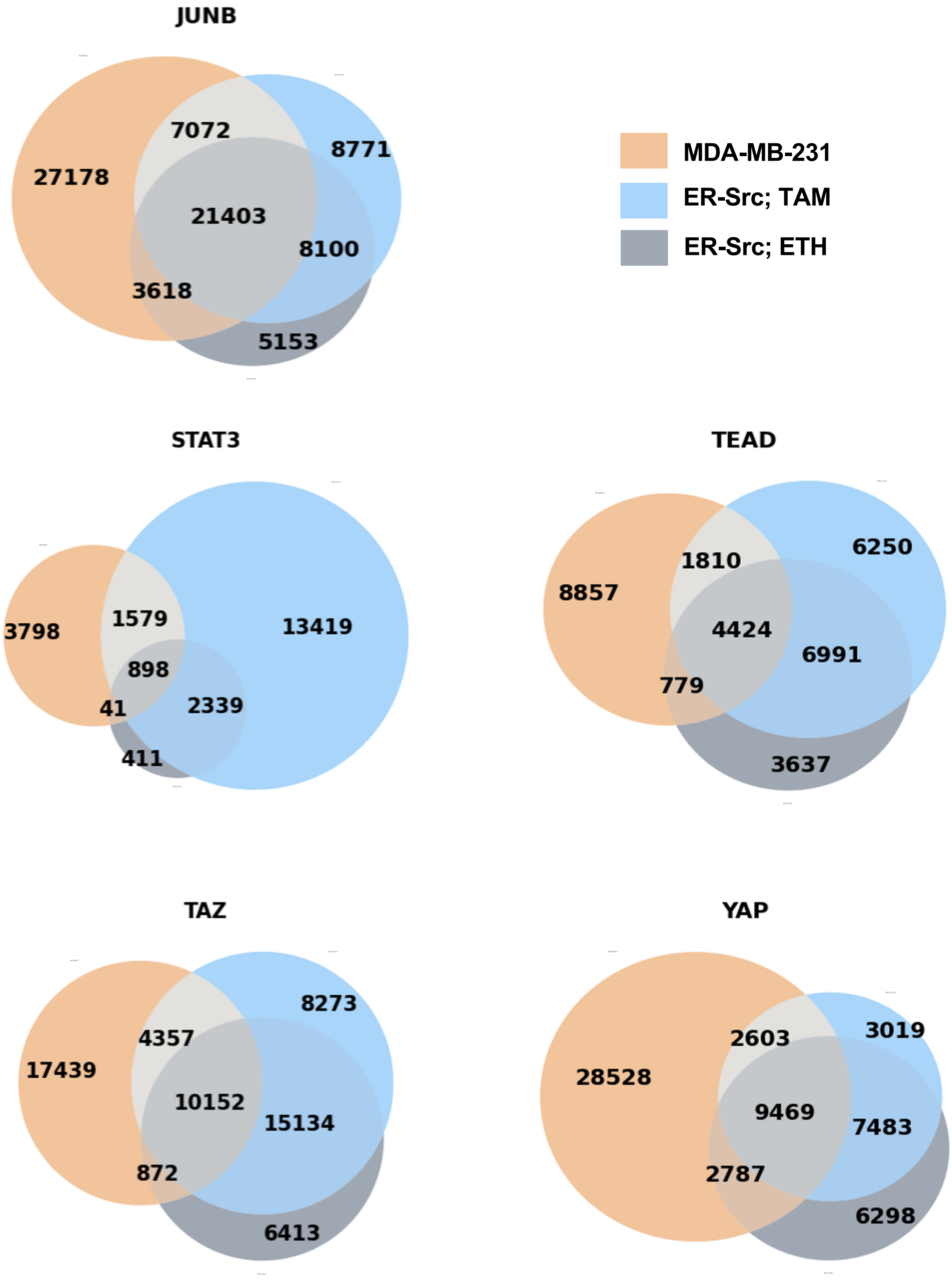

**Figure 6-figure supplement 2**

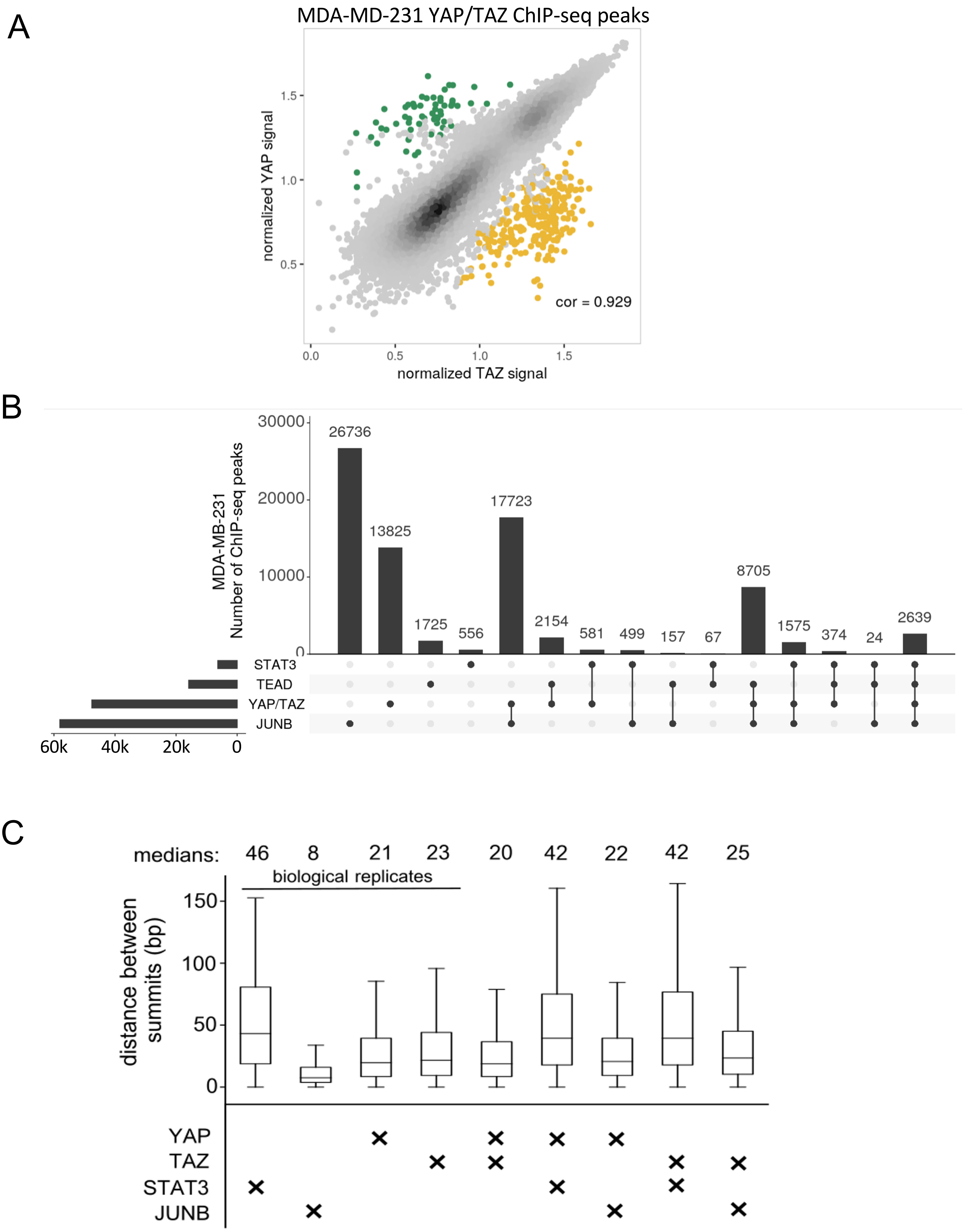

#### Figure 6-figure supplement 3

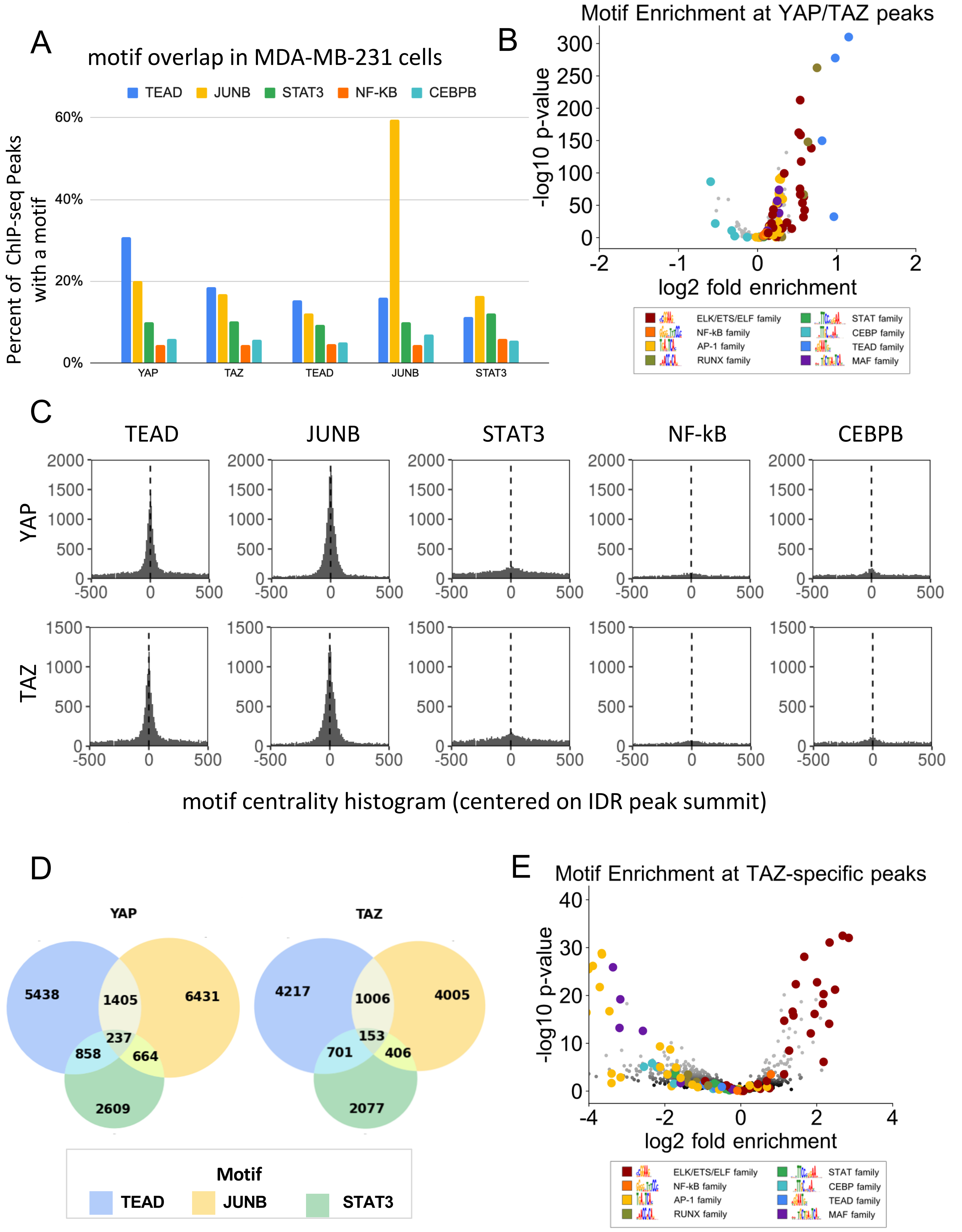

**Figure 6-figure supplement 4**

A

**YAP/TAZ target genes (n=14,851)**

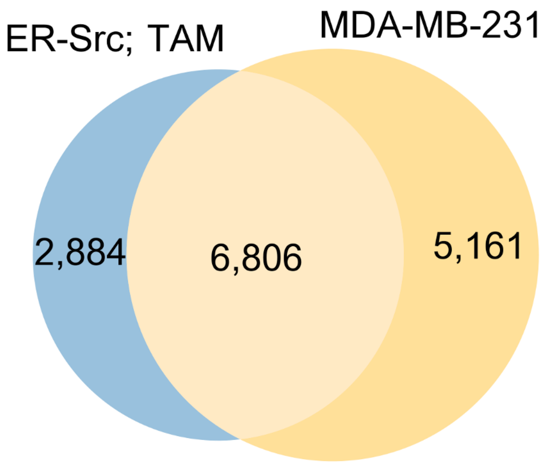


B

**Target genes in MCF10A by ChIP-seq peak (n=14,046)**

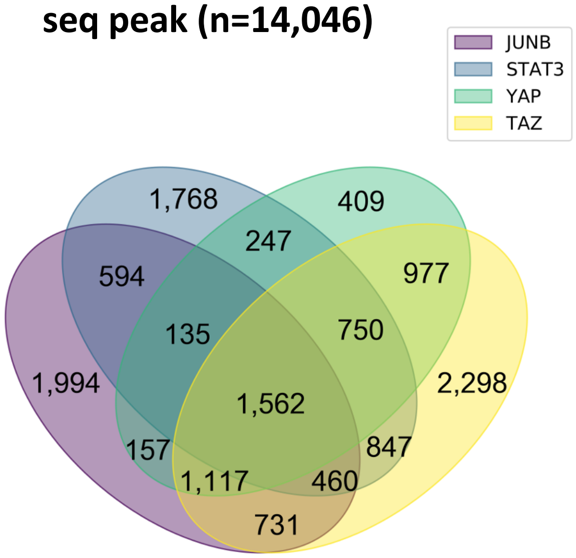


C

**Target genes in MCF10A by motif (n=)**

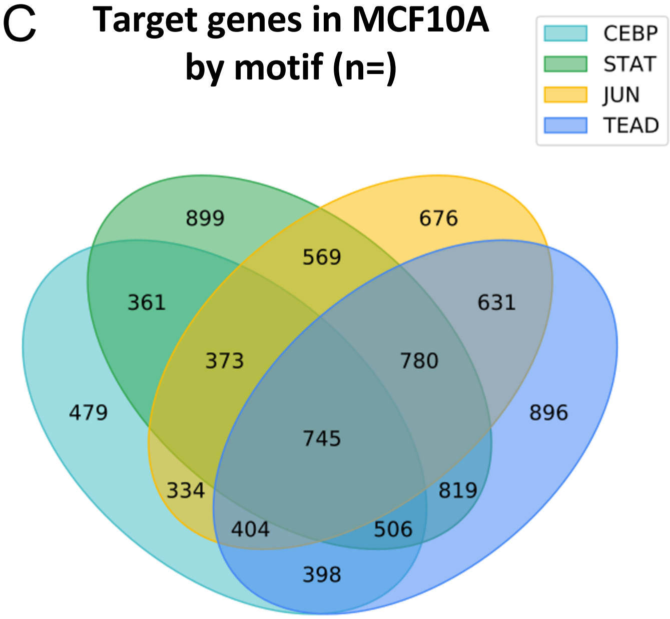


D

**Target genes in MCF10A by motif: GO terms**

**CEBP motif present**

| GO term | p-value | q-value (FDR) |
|---|---|---|
| movement of cell or subcellular component | $3.56\times10^{-7}$ | $4.2\times10^{-3}$ |

**TEAD motif present**

| GO term | p-value | q-value (FDR) |
|---|---|---|
| regulation of cell motility | $1.41\times10^{-6}$ | $8.31\times10^{-3}$ |
| regulation of cellular component movement | $2.1\times10^{-6}$ | $8.29\times10^{-3}$ |

### Figure 7-figure supplement 1

A YAP/TAZ/STAT3/JUNB Gene Signature 1 (533 genes)

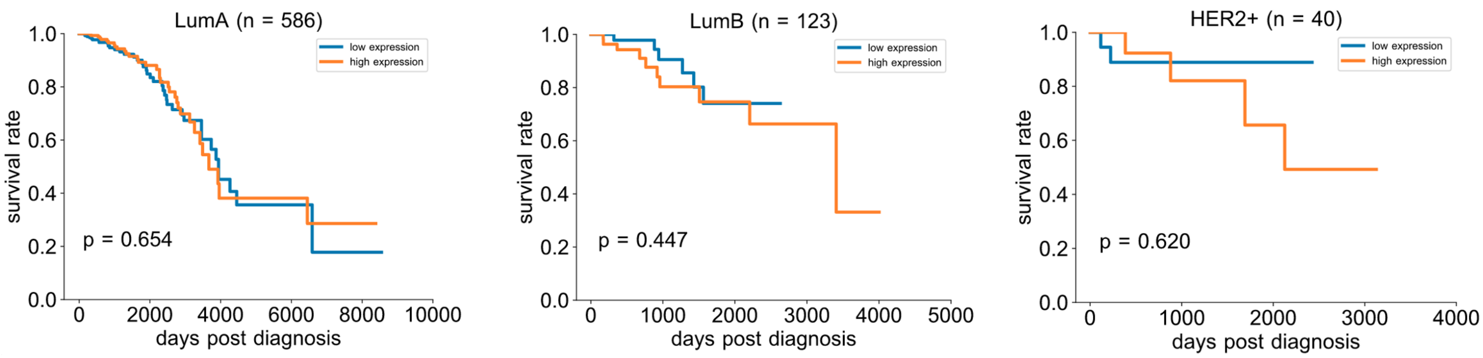


B YAP/TAZ/STAT3/JUNB Gene Signature 2 (563 genes)

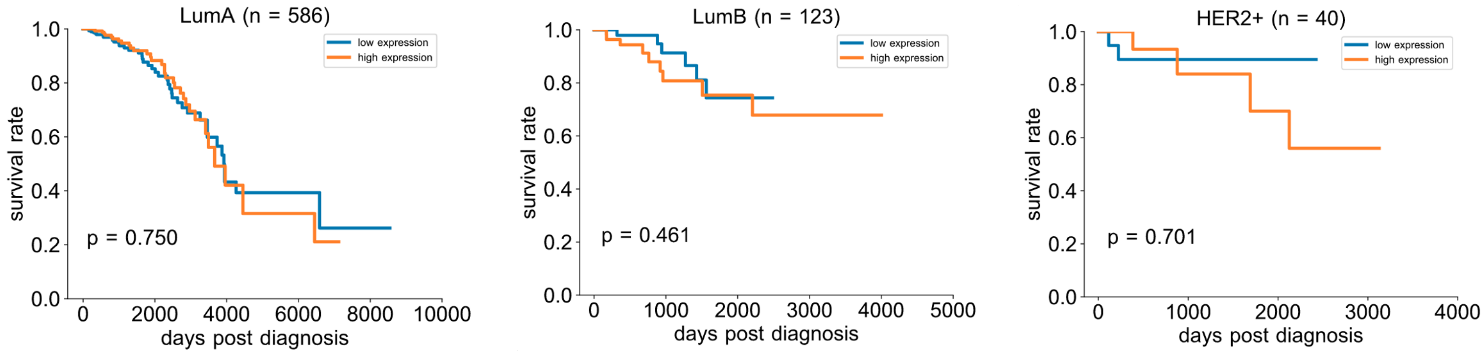


C YAP/TAZ/STAT3/JUNB Gene Signature 3 (45 genes)

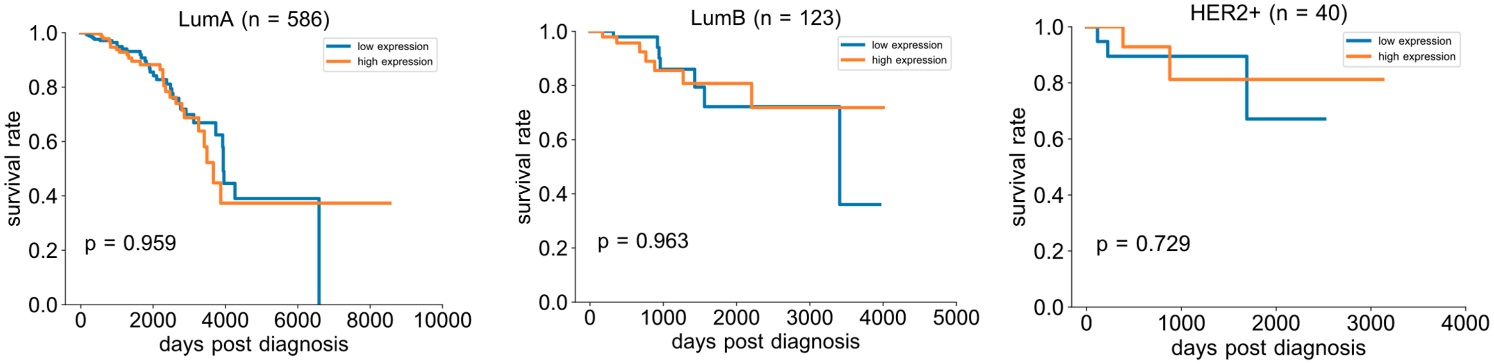


D Sig. 1 random control

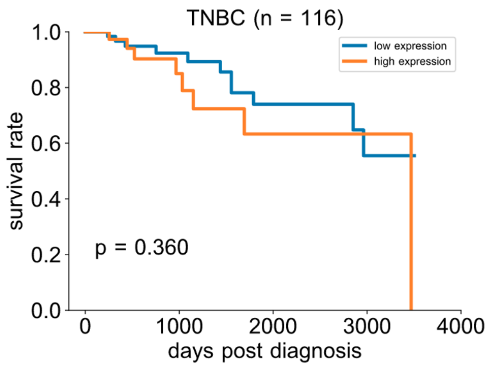


E Sig. 2 random control

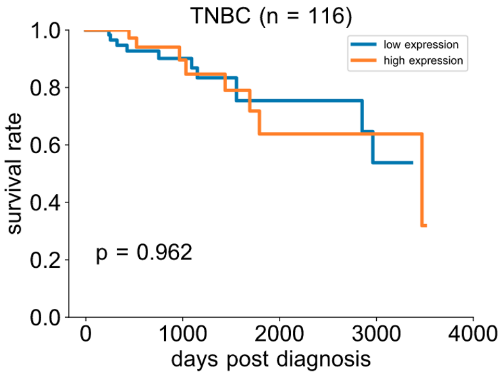


F Sig. 3 random control

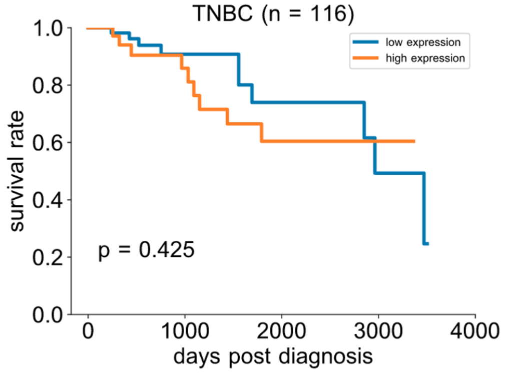

### Figure 7-figure supplement 2

#### A YAP/TAZ/TEAD Gene Signature 1 (533 genes)

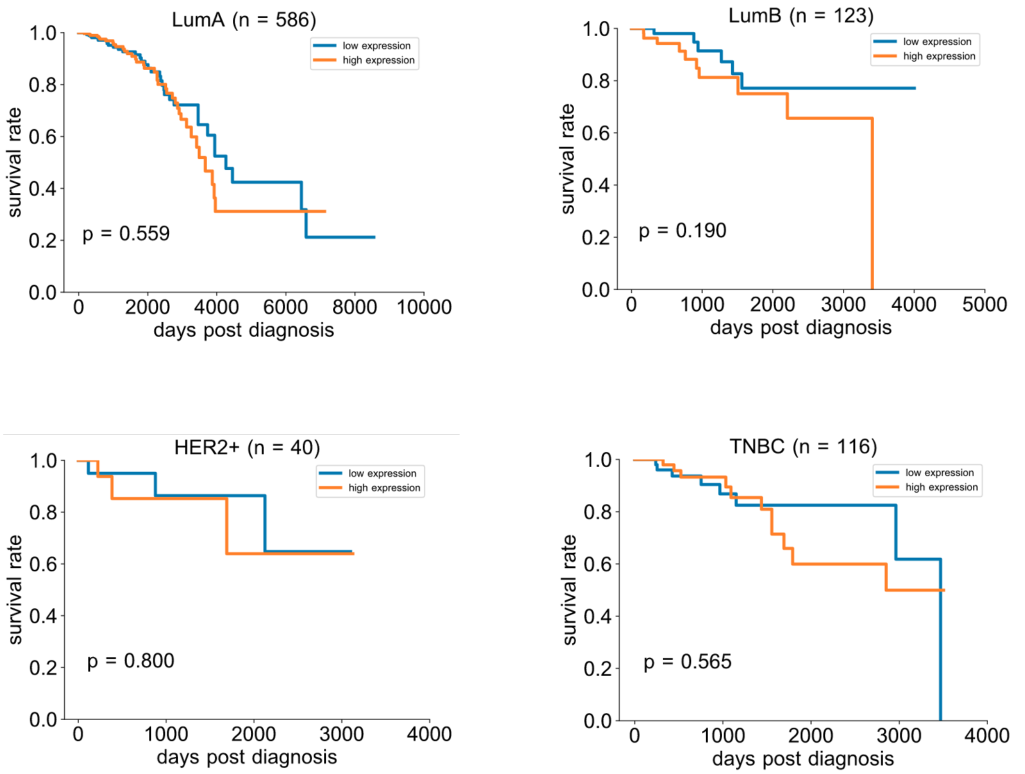


#### B YAP/TAZ/TEAD Gene Signature 2 (52 genes)

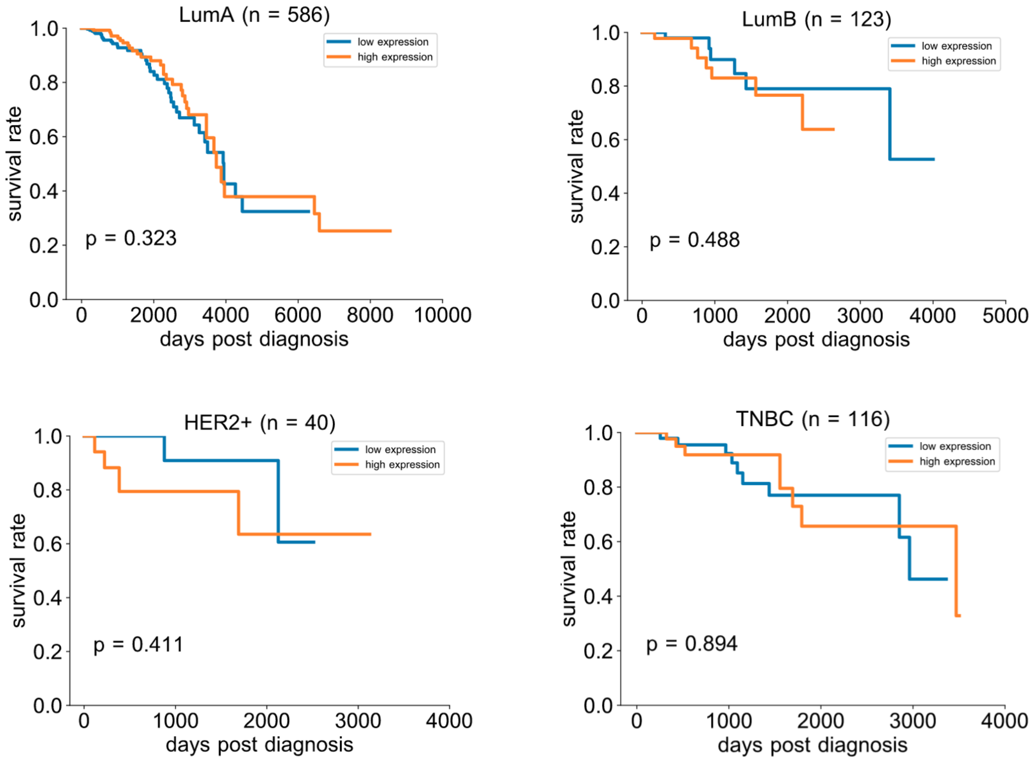
