## Supplementary material for "YAP and TAZ are transcriptional co-activators of AP-1 proteins and STAT3 during breast cellular transformation": Table S1

### A. ChIP-seq signal correlation coefficient between replicates

| TF | YAP | TAZ | TEAD | STAT3 | JUNB |
|---|---|---|---|---|---|
| **ER-Src; ETH** | 0.705 | 0.763 | 0.779 | 0.890 | 0.918 |
| **ER-Src; TAM** | 0.683 | 0.855 | 0.772 | 0.950 | 0.912 |
| **MDA-MB-231** | 0.908 | 0.886 | 0.878 | 0.668 | 0.915 |

### B. ChIP-seq IDR peak numbers

| | YAP | TAZ | TEAD | STAT3 | JUNB |
|---|---|---|---|---|---|
| **ER-Src; ETH** | 25,975 | 32,587 | 15,486 | 3,680 | 37,886 |
| **ER-Src; TAM** | 22,509 | 37,888 | 19,467 | 18,243 | 44,952 |
| **MDA-MB-231** | 43,387 | 32,820 | 15,870 | 6,316 | 59,271 |

| | YAP in TAZ KO | TAZ in YAP KO |
|---|---|---|
| **ER-Src; ETH** | 29,335 | 49,118 |
| **ER-Src; TAM** | 39,230 | 35,221 |
