## Supplementary material for "YAP and TAZ are transcriptional co-activators of AP-1 proteins and STAT3 during breast cellular transformation": Table S2

| GO Term | Description | -log10(q-value), TAZ-specific | -log10(q-value), YAP-specific | -log10(q-value), shared |
|---|---|---|---|---|
| GO:0090304 | nucleic acid metabolic process | 13.64 | 0 | 0 |
| GO:0006139 | nucleobase-containing compound metabolic process | 12.11 | 0 | 0 |
| GO:0034470 | ncRNA processing | 11.92 | 0 | 0 |
| GO:0006396 | RNA processing | 11.79 | 0 | 0 |
| GO:0034660 | ncRNA metabolic process | 11.44 | 0 | 0 |
| GO:0046483 | heterocycle metabolic process | 11.12 | 0 | 0 |
| GO:0016070 | RNA metabolic process | 11.09 | 0 | 0 |
| GO:0034641 | cellular nitrogen compound metabolic process | 10.81 | 0 | 0 |
| GO:0006725 | cellular aromatic compound metabolic process | 10.73 | 0 | 0 |
| GO:1901360 | organic cyclic compound metabolic process | 9.43 | 0 | 0 |
| GO:0006364 | rRNA processing | 6.15 | 0 | 0 |
| GO:0006399 | tRNA metabolic process | 5.96 | 0 | 0 |
| GO:0016072 | rRNA metabolic process | 5.72 | 0 | 0 |
| GO:0044237 | cellular metabolic process | 5.34 | 0 | 0 |
| GO:0070126 | mitochondrial translational termination | 5.04 | 0 | 0 |
| GO:0009451 | RNA modification | 4.49 | 0 | 0 |
| GO:0006415 | translational termination | 4.37 | 0 | 0 |
| GO:0070125 | mitochondrial translational elongation | 3.92 | 0 | 0 |
| GO:0008033 | tRNA processing | 3.85 | 0 | 0 |
| GO:0008152 | metabolic process | 3.82 | 0 | 0 |
| GO:0043624 | cellular protein complex disassembly | 3.32 | 0 | 0 |
| GO:0043170 | macromolecule metabolic process | 3.21 | 0 | 0 |
| GO:0051276 | chromosome organization | 3.18 | 0 | 0 |
| GO:0043933 | protein-containing complex subunit organization | 3.16 | 0 | 0 |
| GO:0019219 | regulation of nucleobase-containing compound metabolic process | 0 | 8.78 | 0 |
| GO:0080090 | regulation of primary metabolic process | 0 | 7.94 | 0 |
| GO:0051171 | regulation of nitrogen compound metabolic process | 0 | 7.87 | 0 |
| GO:0060255 | regulation of macromolecule metabolic process | 0 | 7.8 | 0 |
| GO:0051252 | regulation of RNA metabolic process | 0 | 7.47 | 0 |
| GO:0006357 | regulation of transcription by RNA polymerase II | 0 | 7.25 | 0 |
| GO:2000112 | regulation of cellular macromolecule biosynthetic process | 0 | 7.21 | 0 |
| GO:0010556 | regulation of macromolecule biosynthetic process | 0 | 7.2 | 0 |
| GO:0019222 | regulation of metabolic process | 0 | 7.11 | 0 |
| GO:0050794 | regulation of cellular process | 0 | 7.1 | 0 |
| GO:0031323 | regulation of cellular metabolic process | 0 | 7.1 | 0 |
| GO:0009889 | regulation of biosynthetic process | 0 | 7.09 | 0 |
| GO:1903506 | regulation of nucleic acid-templated transcription | 0 | 7.08 | 0 |

| | | | | |
|---|---|---|---|---|
| GO:0031326 | regulation of cellular biosynthetic process | 0 | 7.06 | 0 |
| GO:2001141 | regulation of RNA biosynthetic process | 0 | 7.03 | 0 |
| GO:0006355 | regulation of transcription, DNA-templated | 0 | 6.63 | 0 |
| GO:0010468 | regulation of gene expression | 0 | 6.45 | 0 |
| GO:0050789 | regulation of biological process | 0 | 6.25 | 0 |
| GO:0051172 | negative regulation of nitrogen compound metabolic process | 0 | 4.96 | 0 |
| GO:0065007 | biological regulation | 0 | 4.55 | 0 |
| GO:0010605 | negative regulation of macromolecule metabolic process | 0 | 3.97 | 0 |
| GO:0031324 | negative regulation of cellular metabolic process | 0 | 3.81 | 0 |
| GO:0006325 | chromatin organization | 0 | 3.58 | 0 |
| GO:1903507 | negative regulation of nucleic acid-templated transcription | 0 | 3.5 | 0 |
| GO:0051253 | negative regulation of RNA metabolic process | 0 | 3.49 | 0 |
| GO:1902679 | negative regulation of RNA biosynthetic process | 0 | 3.48 | 0 |
| GO:0045934 | negative regulation of nucleobase-containing compound metabolic process | 0 | 3.47 | 0 |
| GO:0009892 | negative regulation of metabolic process | 0 | 3.23 | 0 |
| GO:0045935 | positive regulation of nucleobase-containing compound metabolic process | 0 | 3.03 | 0 |
| GO:0010558 | negative regulation of macromolecule biosynthetic process | 0 | 3.02 | 0 |
| GO:0007626 | locomotory behavior | 0 | 0 | 0.53 |
| GO:0032730 | positive regulation of interleukin-1 alpha production | 0 | 0 | 0.33 |
| GO:0032732 | positive regulation of interleukin-1 production | 0 | 0 | 0.29 |
| GO:0000082 | G1/S transition of mitotic cell cycle | 0 | 0 | 0.17 |
| GO:0032650 | regulation of interleukin-1 alpha production | 0 | 0 | 0.15 |
| GO:0032652 | regulation of interleukin-1 production | 0 | 0 | 0.13 |
| GO:0044843 | cell cycle G1/S phase transition | 0 | 0 | 0.12 |
| GO:0006471 | protein ADP-ribosylation | 0 | 0 | 0.09 |
| GO:0048477 | oogenesis | 0 | 0 | 0.04 |
