## Supplementary material for "YAP and TAZ are transcriptional co-activators of AP-1 proteins and STAT3 during breast cellular transformation": Table S3

| GO Term | Description | -log10(q-value), CEBP motif | -log10(q-value), TEAD motif | -log10(q-value), JUN motif | -log10(q-value), STAT motif |
|---|---|---|---|---|---|
| GO:0006954 | inflammatory response | 2.15 | 0 | 0 | 0 |
| GO:0048878 | chemical homeostasis | 1.56 | 0 | 0 | 0 |
| GO:0030029 | actin filament-based process | 0 | 3.33 | 0 | 0 |
| GO:0030036 | actin cytoskeleton organization | 0 | 2.46 | 0 | 0 |
| GO:0007165 | signal transduction | 0.56 | 2.38 | 1.38 | 0 |
| GO:0032879 | regulation of localization | 0.65 | 1.13 | 0.97 | 0 |
| GO:0097435 | supramolecular fiber organization | 0 | 1.08 | 0 | 0 |
| GO:0032501 | multicellular organismal process | 0 | 1.07 | 0.77 | 0 |
| GO:0007155 | cell adhesion | 0 | 1.05 | 1.63 | 0 |
| GO:0022610 | biological adhesion | 0 | 1.03 | 1.64 | 0 |
| GO:0031032 | actomyosin structure organization | 0 | 1 | 0 | 0 |
| GO:0032502 | developmental process | 0 | 0 | 3.47 | 0 |
| GO:0048856 | anatomical structure development | 0 | 0 | 2.91 | 0 |
| GO:0051239 | regulation of multicellular organismal process | 0 | 0 | 2.03 | 0 |
| GO:0065007 | biological regulation | 0 | 0.48 | 1.96 | 0 |
| GO:2000026 | regulation of multicellular organismal development | 0 | 0 | 1.64 | 0 |
| GO:0007229 | integrin-mediated signaling pathway | 0 | 0 | 1.56 | 0 |
| GO:0009888 | tissue development | 0 | 0 | 1.54 | 0 |
| GO:0048646 | anatomical structure formation involved in morphoge | 0 | 0 | 1.54 | 0 |
| GO:0023051 | regulation of signaling | 0 | 0 | 1.52 | 0 |
| GO:0010646 | regulation of cell communication | 0 | 0 | 1.51 | 0 |
| GO:0043062 | extracellular structure organization | 0 | 0 | 1.39 | 0 |
| GO:0050789 | regulation of biological process | 0 | 0.43 | 1.27 | 0 |
| GO:0048870 | cell motility | 0 | 0.46 | 1.21 | 0 |
| GO:0031581 | hemidesmosome assembly | 0 | 0 | 1.19 | 0 |
| GO:0030198 | extracellular matrix organization | 0 | 0 | 1.19 | 0 |
| GO:0048869 | cellular developmental process | 0 | 0 | 1.19 | 0 |
| GO:0010631 | epithelial cell migration | 0 | 0 | 1.17 | 0 |
| GO:0002576 | platelet degranulation | 0 | 0 | 1.1 | 0 |
| GO:0043542 | endothelial cell migration | 0 | 0 | 1.08 | 0 |
| GO:0009966 | regulation of signal transduction | 0 | 0 | 1.08 | 0 |
| GO:0050793 | regulation of developmental process | 0 | 0 | 1.08 | 0 |
